# Systems-Level Proteomics Evaluation of Microglia Response to Tumor-Supportive Anti-inflammatory Cytokines

**DOI:** 10.1101/2021.02.04.429830

**Authors:** Shreya Ahuja, Iulia M. Lazar

## Abstract

**Background:** Microglia safeguard the CNS against injuries and pathogens by inducing an inflammatory response. When exposed to anti-inflammatory cytokines, these cells possess the ability to switch from an inflammatory to an immunosuppressive phenotype. Cancer cells exploit this property to evade the immune system, and elicit an anti-inflammatory microenvironment that facilitates tumor attachment and growth.

**Objective:** The tumor-supportive biological processes that are activated in microglia cells in response to anti-inflammatory cytokines released from cancer cells were explored with mass spectrometry and proteomic technologies.

**Methods:** Serum-depleted and non-depleted human microglia cells (HMC3) were treated with a cocktail of IL-4, IL-13, IL-10, TGFβ, and CCL2. The cellular protein extracts were analyzed by LC-MS/MS. Using functional annotation clustering tools, statistically significant proteins that displayed a change in abundance between cytokine-treated and non-treated cells were mapped to their biological networks and pathways.

**Results:** The proteomic analysis of HMC3 cells enabled the identification of ∼10,000 proteins. Stimulation with anti-inflammatory cytokines resulted in the activation of distinct, yet integrated clusters of proteins that trigger downstream a number of tumor-promoting biological processes. The observed changes could be classified into four major categories, i.e., mitochondrial gene expression, ECM remodeling, immune response, and impaired cell cycle progression. Intracellular immune activation was mediated mainly by the transducers of MAPK, STAT, TGFβ, NFKB, and integrin signaling pathways. Abundant collagen formation along with the expression of additional receptors, matrix components, growth factors, proteases and protease inhibitors, enabled ECM remodeling processes supportive of cell-cell and cell-matrix adhesion. Overexpression of integrins and their modulators was reflective of signaling processes that correlated ECM reorganization with cytoskeletal re-arrangements supportive of cell migration. Antigen processing/presentation was represented by HLA class I histocompatibility antigens, and correlated with upregulated proteasomal subunits, and vesicular/viral transport and secretory processes. Immunosuppressive and proangiogenic chemokines were detectable in low abundance. Pronounced pro-inflammatory, chemotactic or phagocytic trends were not observed, however, the expression of certain receptors and ECM proteins indicated the presence of such capabilities.

**Conclusions:** Comprehensive proteomic profiling of HMC3 cells stimulated with anti-inflammatory cytokines revealed a microglia phenotype that provides novel insights into the tumor microenvironment-driven mechanisms that fuel cancer development in the brain.

## Introduction

The macrophage glial network population of the central nervous system (CNS), also called “microglia,” was first studied in 1919 by a Spanish researcher, Pio del Rio-Hortega who defined these entities as “voracious macrophages” of the brain parenchyma. Morphologically, microglia cells are found in the brain in a ramified form, i.e., having a small cell body with long branched protrusions which the cells constantly extend and retract to inspect the environment for toxic or necrotic molecules, or for pathogens. In this way, the cells act as the surveillance system of the CNS, constantly patrolling the region and responding by activating a repertoire of cytotoxic factors that destroy the invaders [1,2]. Once a damage is detected, they rapidly transform into “reactive” ameboid mobile effector cells that can proliferate, phagocytose and interact with other immune cells via antigen presentation. Therefore, microglia play a crucial role in maintaining and restoring the homeostasis of the CNS. While serving as immune guardians, microglia have the ability to engage in distinct response programs which may be cytotoxic or immunosuppressive, with implications for CNS infections and cancer [3]. Microglia cells exhibit dual behavior, one that recognizes and attacks injuries, infections and damages to the CNS, i.e., the “classical” M1 phenotype; and, a second one that dampens inflammatory responses to support for example tumor progression, i.e., the “alternative” M2 phenotype [4]. The polarization of microglia toward either phenotype is triggered by a specific concoction of cytokines. Stimulation toward the M1 phenotype occurs in response to bacterial endotoxin lipopolysaccharides (LPS) or interferons (IFN-γ), which render the microglia bactericidal and immunostimulant. In this subtype, there is an increased expression of STAT1 in the cells that further upregulates the expression of pro-inflammatory cytokines (IL-1β, IL-12, TNF-α) and of oxidative metabolites (iNOS) [5]. On the contrary, the M2 phenotype is elicited when tumor cells in the brain “hijack” the pro-inflammatory nature of the microglia by sending out signals that promote tumor progression through processes such as matrix deposition and tissue remodeling [6]. Chemoattractants such as CCL2 (or MCP1), secreted by cancer cells, can stimulate microglia via CCR2 activation, and recruit the microglia to the tumor microenvironment to support invasiveness through IL-6 expression [7]. IL-6 activates an “M2” specific enzyme, Arginase-1 (ARG1) [8], that can alter the extracellular matrix composition to drive cancer progression. Additionally, anti-inflammatory cytokines such as IL-10, IL-13, IL-4 and TGFβ skew the microglia toward an M2 phenotype by increasing the intracellular levels of STAT3, which then negatively affects the expression of MHC-II, CD80, CD86 [5,9-11], and upregulates the expression of the macrophage scavenger receptor 1 (MSR1)/CD204 and of CD163 [5].

The brain offers a privileged, immunocompetent niche which is safeguarded from insults to the rest of the body. The blood brain barrier isolates the brain from the outside environment [12], thereby offering the perfect lodging spot for circulating cancer cells. Therefore, the brain is a “popular” destination for metastasized cancers which even surpass the incidents of primary brain tumors. Carcinomas such as those of the breast, kidney, lung and melanoma are the most frequent to metastasize to the brain [13], with a median survival rate ranging from 6 to 12 months, depending on the type and grade of primary tumor and extent of the systemic disease [14]. Numerous reports have suggested that resident brain cells such as microglia support the survival and growth of metastasized cancer due to their ability to respond favorably to signals from the tumor microenvironment [15]. Despite the prominence of these immune sentinels in the CNS and their role in the establishment of brain metastases, the interactions of microglia with the tumor cells and the acquisition of distinct anti-inflammatory properties is yet to be fully explored. The field is primarily limited by the inability to differentiate the brain microglia from the infiltrated bone-marrow derived macrophages [16]. Many groups have reported similarities between the M2 activation of microglia and of tumor-associated macrophages (TAMs) that contribute to an immunosuppressive and tumor supporting milieu, however, the exact mechanisms of microglial plasticity, especially their re-programming during cancer metastasis, is still ambiguous to the scientific community [17]. Moreover, due to ethical difficulties associated with the procurement of brain samples or isolation of primary cell cultures, studies focused on clarifying the role of microglia in the establishment of metastasized cancer have been mostly performed using rodent models [18,19]. Recently, however, the introduction of human microglia cell lines has been pivotal in overcoming these issues. Primary human microglia cells can be transfected with bacterial plasmids encoding for the SV40 large T-antigen to generate continuous cell lines [20,21]. In contrast to primary cells which have low renewing efficiency, such cell lines offer the ease of *in-vitro* culture and yield adequate amounts for sample analysis. Easy upscaling is also supportive of large-scale omics projects, as demonstrated in a study in which IFN-γ and IL-4 were used to induce M1 or M2 polarization, respectively, to investigate the proteomic differences between the two phenotypes [22]. Comprehensive characterization of the M1/M2 microglia activation states and of their role in brain metastases has not been performed, however, and represents therefore the need of the hour. To address the paucity of information in this field and advance the understanding of metastatic cancer cell development in the brain, in this study, we used mass spectrometry (MS) and proteomics to extensively characterize the human fetal brain-derived microglial cell line (HMC3) under basal and anti-inflammatory cytokine stimulation conditions. We explored the data for the presence of microglia-specific markers and of cell surface proteins with immunological functions, and for quantitative changes in protein expression between cytokine-treated and non-treated cells. The results were assessed and discussed in terms of altered expression of specific sets of proteins and biological processes that are crucial for the suppression of immune activation and the formation of a tumor-supportive niche.

## Experimental methods

### Materials

HMC3 cells, Eagle’s minimum essential medium and penicillin-streptomycin (PenStrep) solution were purchased from ATCC (Manassas, VA). Phenol red and glutamine free Minimum Essential Medium (MEM), L-glutamine, Dulbecco’s phosphate buffered saline (DPBS) and trypsin-EDTA were purchased from Gibco (Gaithersburg, MD). Fetal bovine serum (FBS) was supplied by Gemini Bio Products (West Sacramento, CA). Recombinant human IL-4, IL-10, IL-13, CCL2, TGFβ1 (HEK293 derived) and TGFβ2 (HEK293 derived) were acquired from Peprotech (Rocky Hill, NJ). Primary antibody [mouse monoclonal IgG anti-Vimentin (V9)], secondary antibody mouse IgG conjugated to CruzFluor488 (sc-516176), UltraCruz® blocking reagent, and UltraCruz® aqueous mounting medium with DAPI were purchased from Santa Cruz Biotechnology (Dallas, TX). Propidium iodide for FACS analysis was from Invitrogen (Carlsbad, CA). Cell Lytic™ NuCLEAR™ extraction kit for the separation of nuclear and cytoplasmic fractions of cells, trifluoroacetic acid (TFA), protein standards, phosphatase inhibitors [sodium orthovanadate (Na_3_VO_4_) and sodium fluoride (NaF)], ammonium bicarbonate (NH_4_HCO_3_), urea, dithiothreitol (DTT), ribonuclease (RNase) and Triton X-100 were procured from Sigma Aldrich (St. Louis, MO). Bradford reagent and bovine serum albumin (BSA) standards were provided by Biorad (Hercules, CA). Trypsin, sequencing grade, was from Promega (Madison, WI). Sample clean-up SPEC-PTC18 and SPEC-PTSCX pipette tips were bought from Agilent technologies (Santa Clara, CA). HPLC grade acetonitrile and methanol were from Fischer Scientific (Fair Lawn, NJ). Ethanol was obtained from Decon Laboratories (King of Prussia, PA). High purity water was prepared in-house by distillation of de-ionized water.

### Cell culture

HMC3 cells were authenticated by short-tandem repeat (STR) profiling at ATCC. Three batches of cells, revived from liquid nitrogen, were used to produce three biological replicates of each experimental cell treatment. HMC3 cells were routinely cultured in EMEM supplemented with FBS (10 %), in a water jacketed incubator at 37 °C and with 5 % CO_2_. PenStrep (0.5 %) was added to all culture media. For generating control cultures, the cells were either starved for 48 h in EMEM, or starved for 48 h and then released with EMEM supplemented with FBS (10 %) for 24 h. For cytokine-stimulated cultures, the HMC3 cells were starved for 24 h in phenol red-free MEM with glutamine (2 mM), and then for another 24 h with MEM supplemented with a cytokine cocktail. Alternatively, cells starved for 48 h in EMEM were stimulated with EMEM supplemented with FBS (10 %) and cytokines. The cytokines were added in concentrations that were proven effective in previously published works [22-24]. The cytokine cocktail included TGFβ1 (20 ng/mL), TGFβ2 (20 ng/mL), IL-13 (20 ng/mL), IL-10 (20 ng/mL), IL-4 (40 ng/mL), and CCL2 (40 ng/mL).

### FACS analysis

The cells were fixed using 70 % ethanol and then stained with propidium iodide (0.02 mg/mL) in a PBS solution containing Triton X-100 (0.1 %) and RNAase (0.2 mg/mL). The cells were incubated for 30 min at room temperature in the staining solution before being subjected to flow cytometry analysis (FACSCalibur, BD Biosciences, San Jose, CA).

### Immunofluorescence microscopy

Immunofluorescent detection of vimentin was performed by fixing the cells with chilled methanol (-20 °C, 15 min), blocking (1 h), incubating with primary antibody (2 ug/mL in blocking buffer, 1 h), and staining with the secondary antibody conjugated to the detection fluorophore (1 ug/mL in blocking buffer, 30 min). The cells were visualized with an inverted Eclipse TE2000-U epi-fluorescence microscope (Nikon Instruments Inc., Melville, NY). Images were acquired and processed with a Nikon’s NIS-Elements Advanced Research imaging platform, version 5.11.01.

### Sample preparation for MS analysis

The cells were harvested by trypsinization at the end of each treatment and flash frozen at -80 °C until further use. Separation of nuclear and cytoplasmic fractions was carried out in accordance with the protocol described in the Cell Lytic™ NuCLEAR™ extraction kit. The lysis reagents were supplemented with phosphatase and protease inhibitors. A Bradford assay was performed to measure the concentration of the protein extracts. Protein extracts (500 ug) were denatured and reduced with 8 M urea and 5 mM DTT for 1 h at 57 °C. The reduced proteins were digested overnight with trypsin at 50:1 protein-to-enzyme ratio at 37 °C. The reaction was quenched with glacial acetic acid, and the samples were subjected to C18/SCX clean-up. The protein digests were re-suspended in H_2_O:CH_3_CN:TFA (98:2:0.01) at a concentration of 2 μg/μL [25,26], and frozen at -80 °C until liquid chromatography (LC)-MS analysis was conducted.

### LC-MS analysis

LC separations were performed with an EASY-nLC 1200 UHPLC system by using an EASY-Spray column ES802A (250 mm long, 75 μm i.d., 2 μm C18/silica particles, ThermoFisher Scientific) operated at 250 nL/min. Mobile phase A consisted of 0.01 % TFA in H_2_O:CH_3_CN (96:4, v/v), and mobile phase B of 0.01 % TFA in H_2_O:CH_3_CN (10:90, v/v). The solvent gradient was ∼2 h long, with increasing concentration of solvent B according to the following steps: 7 % B, 2 min; 7-30 % B, 105 min; 30-45 % B, 2 min; 45-60 % B, 1 min; 60-90 % B, 1 min; 90 % B, 10 min; 90-5 % B, 1 min; 7 % B, 5 min. Mass spectrometry analysis was performed with a Q Exactive hybrid quadrupole-Orbitrap mass spectrometer (ThermoFisher Scientific). Nano-electrospray ionization (ESI) was induced at 2 kV, and the ion transfer capillary temperature was set to 250 °C. Full mass spectra were acquired over a scan range of 400-1600 m/z, with acquisition parameters set to resolution 70,000, AGC target 3E6, and maximum IT 100 ms. The quadrupole isolation width was 2.4 m/z, and collision induced dissociation (CID) was performed at a normalized collision energy of 30 %. Data dependent MS2 acquisition was completed with a resolution setting of 17,500, AGC target 1E5, maximum IT 50 ms, and loop count of 20. Other data dependent settings included minimum AGC target 2E3 (with resulting intensity threshold of 4E4), apex trigger 1 to 2 s, charge exclusion enabled for unassigned and +1 charges, isotope exclusion on, preferred peptide match on, and dynamic exclusion of 10 s for chromatographic peak widths of 8 s.

### MS data processing

Peptide-spectrum matches (PSM) and protein identifications were performed with the Sequest HT search engine embedded in the Proteome Discoverer (v. 2.4) software package (Thermo Fisher Scientific, Waltham, MA), and a target/decoy-based processing workflow of peptide precursor masses between 400 and 5,000 Da. The Sequest HT node searches were made against a *Homo sapiens* database, downloaded from UniProtKB (March 2019), containing 20,404 reviewed/non-redundant sequences. The workflow parameters were set up for the identification of only fully tryptic peptides with a min of 6 and max of 144 amino acids, max 2 missed cleavages, precursor ion tolerance of 15 ppm, and fragment ion tolerance of 0.02 Da. All **b/y/a** ion fragments were considered, with dynamic modifications on Met (oxidation) and the N-terminal amino acid (acetyl). The signal-to-noise (S/N) ratio threshold was set to 1.5. The target/decoy PSM validator node included concatenated databases, with input data filtered for maximum DeltaCn of 0.05 and maximum rank of 1, and strict/relaxed target PSM FDRs of 0.01 and 0.03, respectively. In the consensus workflow, the peptide group modification site probability threshold was set to 75, and the peptide validator node to automatic mode with strict/relaxed PSM and peptide target FDRs of 0.01/0.03 (high/medium). The peptide/protein filter node retained only peptides with at least medium confidence that were min 6 amino acids in length, and only proteins matched by rank 1 peptides. Peptides were counted only for the top scoring proteins. The strict/relaxed confidence thresholds for the protein FDR validator node were set again to 0.01/0.03, and the strict parsimony principle was enabled for the protein grouping node.

### Statistical and bioinformatics analysis of MS data

For each of the 24 biological samples (cytokine-treated and control cells, serum-deprived and non-deprived, nuclear and cytoplasmic fractions, three biological replicates of each), three LC-MS/MS technical replicates were generated and combined during the database search process to produce one multiconsensus search file. A global multi-consensus report, compiled from the 72 MS-MS experiments performed for the 24 *samples, was then generated by using Proteome Discoverer 2*.*4 and the results were aligned with* the *Homo sapiens* database IDs for downstream analysis. Fold-changes (FC) for differentially expressed proteins between cytokine-treated and non-treated cells were assessed based on spectral count data, and statistical significance was calculated by performing a t-test. For each comparison, data normalization was performed based on the average of total spectral counts of the six samples taken into consideration, i.e., of the three control and three cytokine-treated biological cell replicates. The resultant normalization factor was used to normalize the counts of each individual protein in the six samples. Missing values in the proteomics data (i.e., proteins with zero counts) were handled by adding a spectral count of 1 to each protein. Differentially expressed proteins were then derived by calculating the Log_2_(Treatment/Control) spectral count ratios, and computing a two-tailed t-test for each protein. Proteins matched by two distinct peptide sequences with a 2-FC in spectral counts, i.e., Log_2_(Treatment/Control)≥0.9 or ≤(-0.9) with a p-value<0.05, were selected for differential expression analysis. These proteins were then interpreted in a biological context using bioinformatics tools provided by online data processing packages such as DAVID 6.8 (*www.david.ncifcrf.gov*) [27], STRING 11.0 *(www.string-db.org*) [28], KEGG (*https://www.genome.jp/kegg/*) [29] and Reactome *(https://reactome.org*) [30]. Functionally related protein lists were created based on controlled vocabulary terms from UniProt (*https://www.uniprot.org/*) [31], and protein functionality was extracted either from the UniProt or GeneCards (*https://www.genecards.org/*) [32] databases. Biological pathways were created with BioRender *(https://biorender.com/*) [33].

## Results

### Rationale

The routine function of microglia, which is to protect the CNS, is disrupted in the event of brain cancer. Glioblastoma and brain metastasis model systems have shown that the immune response of microglia cells is greatly compromised in the presence of cancer cells that engage in a cross-talk with microglia to induce the growth and survival of the former. This occurs in response to anti-inflammatory cytokines released from cancer cells that promote the development of tumor-supportive/pro-neoplastic microglia in a process known as M2 polarization [34]. In this study, we used MS-based proteomics to examine HMC3 activation in response to stimulation with a specific set of molecular factors known to originate from cancer cells during brain metastasis. We hypothesized that the HMC3 cells, upon stimulation with anti-inflammatory cytokines, will show an increased expression of immune-suppressive and tumor-supportive proteins and associated pathways. A cytokine cocktail composed of interleukins IL-4, IL-13, IL-10, growth factors TGFβ1/TGFβ2, and chemokine CCL2 was used for this purpose. These factors have been extensively studied in various animal and macrophage model systems, but much less in human microglia, and found to be key players in promoting tumorigenesis in the brain [5,35-37]. TGFβ is one of the most important anti-inflammatory cytokines that augments the action of IL-4 in promoting an M2 pro-tumor phenotype in a MAPK dependent signaling manner [38]. IL-4 works in concert with IL-13 and IL-10 to execute anti-inflammatory roles that limit the production of inflammatory chemokines by downregulating the activity of STAT1 and NFKβ [39]. Cell activation by IL-4 and IL-13 has been also shown to downregulate the expression of certain ECM remodeling matrix metalloproteinases (MMPs) [40], and to upregulate Arginase-1 [41] to lay the groundwork for collagen formation, tissue repair and cell growth. MCP1/CCL2, a chemokine, helps in recruiting monocytes to the site of tumorigenesis [42]. The decision for stimulating the HMC3 cells with a combination of cytokines was built on two premises, i.e., (a) *in vivo*, the crosstalk between tumor cells and microglia is supported by an exchange of multiple cytokines rather than just one, and (b) if used alone, none of the cytokines will result in an amplified response in microglial cells. This approach was backed by a previous study [43] that reported that CCL2 singlehandedly was unable to alter microglial function in culture, and that other factors might be needed to induce an observable effect.

### Experimental design and observations

As resident immune cells of the brain, microglia cells have an extremely slow turnover rate throughout adulthood, making them one of the slowest dividing cells of the immune system [44]. The “resting” and “activated” states of microglia were mimicked with HMC3 cells cultured in the absence and presence of FBS and cytokines, respectively. Serum deprivation was also used to provide a basal cell state level for enabling a more accurate observation of cytokine-stimulated behavior without interference from growth promoting factors from serum. Cells grown in culture media deprived of FBS for 48 h, or deprived of FBS for 48 h and then treated with FBS for 24 h, were used as control. Serum deprivation was expected to generate a G1 cell cycle arrest-like state, while culture in the presence of FBS, a state typical to cells in the S-stage of the cell cycle. Stimulation with anti-inflammatory cytokines was performed for both cell states. For G1-cells, 24 h of serum deprivation was followed by 24 h stimulation with cytokines. For S-released cells, the cytokines were added together with FBS for 24 h after 48 h serum deprivation. Three biological replicates were generated for each control and stimulated cells, and cellular fractionation into nuclear (N) and cytoplasmic (C) lysates was performed to allow for the generation and characterization of an enriched pool of nuclear proteins. To increase the number of identified proteins, the confidence in protein identifications, and the consistency and reproducibility of proteomic profiling results among biological replicates, three LC-MS/MS technical replicates were acquired for each cell state. The experimental setup along with sample annotations are depicted in **Figure 1A**. Serum-deprived cells were denoted as G1 states, non-deprived cells as S states, and the cytokine-treated cells by a subscript “ck.”

**Figure 1.**
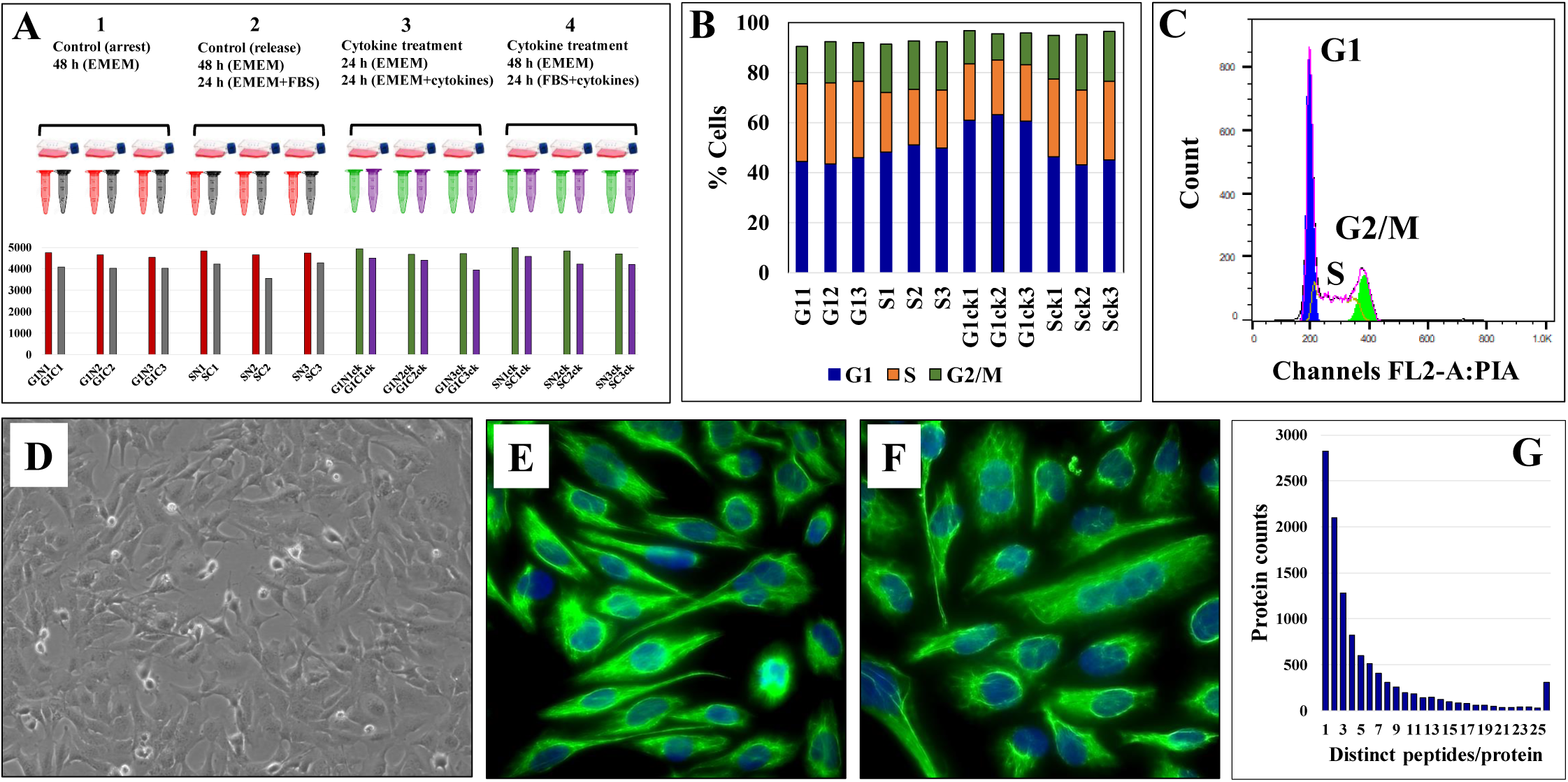
Evaluation of HMC3 cell culture protocol. (**A**) Experimental setup for cell analysis. Three biological cell replicates were cultured in serum-rich and serum-free media, in the presence and absence of cytokines. Cells were fractionated in nuclear and cytoplasmic fractions and analyzed by L-MS/MS to yield ∼4,000 protein IDs per 3 replicate/combined injections. (**B**) Stacked bar graph of HMC3 cell cycle distributions in the various experimental cell states. (**C**) Representative FACS analysis plot of HMC3 cells: G1 45-50 %, S 22-30 %, G2/M 10-20 %. (**D**) Phase contrast microscopy image of proliferating HMC3 cells (20X magnification). (**E**) Immunofluorescence image of resting HMC3 cells stained for vimentin cytoskeletal filaments with XNAMEX primary antibody and secondary mouse IgG antibody conjugated to CruzFluor488 (green fluorescence); nuclei stained with DAPI (blue fluorescence). (**F**) Immunofluorescence image of activated HMC3 cells (same conditions as in D). (**G**) Histogram of protein counts distributed based on the number of matching unique peptides.

FACS analysis revealed that the proportion of G1 and S cells did not change significantly during the different treatment regimens (∼45-50 % cells in G1 and ∼22-30 % in S; CV=3-11%; **Figures 1B** and **C**). Major morphological differences between serum-deprived and non-deprived cells were not observed either, fluorescently labeled cells for vimentin - a cytoskeletal mesenchymal type III intermediate filament protein in microglia - confirming the same cell morphology for the two cell states [45] **(Figures 1D-F)**. As microglia cells were shown to upregulate autophagic behavior in response to serum starvation [46], the lack of cell cycle arrest was not unexpected. Oncogenic transformation as a result of primary cell transfection could have been an additional contributing factor [47-49]. Nevertheless, cytokine-treated cells in serum-deprived medium exhibited a larger proportion of G1 cells (∼60 %, **Figure 1B**), which, as will be discussed later, was an expected result of the addition of cytostatic TGFβ to the cell culture [50].

### Data analysis

The combined analysis of cytokine-treated (G1ck, Sck) and untreated (G1, S) HMC3 cells led to the identification of a total of 10,832 high (1 % FDR) and medium (3 % FDR) confidence protein groups, with an average of 4,463 (CV=8 %) protein groups per each G1 or S cell state and nuclear or cytoplasmic fraction, and 6,136 (CV=7.6 %) protein groups for three combined biological replicates of each cell state (**Figure 1A** and **Supplemental Table 1**). **Supplemental Table 2** provides a description of the gene abbreviations used throughout this study. As many as 75 % of the proteins were matched by two or more distinct peptides (**Figure 1G**). Scatterplot correlation diagrams between the peptide spectrum matches (PSMs) of a protein in two biological replicates (Pearson correlation coefficient R>0.98), and Venn diagrams of protein overlaps in three biological replicates (∼50 % overlap), illustrate the reproducibility of protein identifications (**Figure 2**).

**Figure 2.**
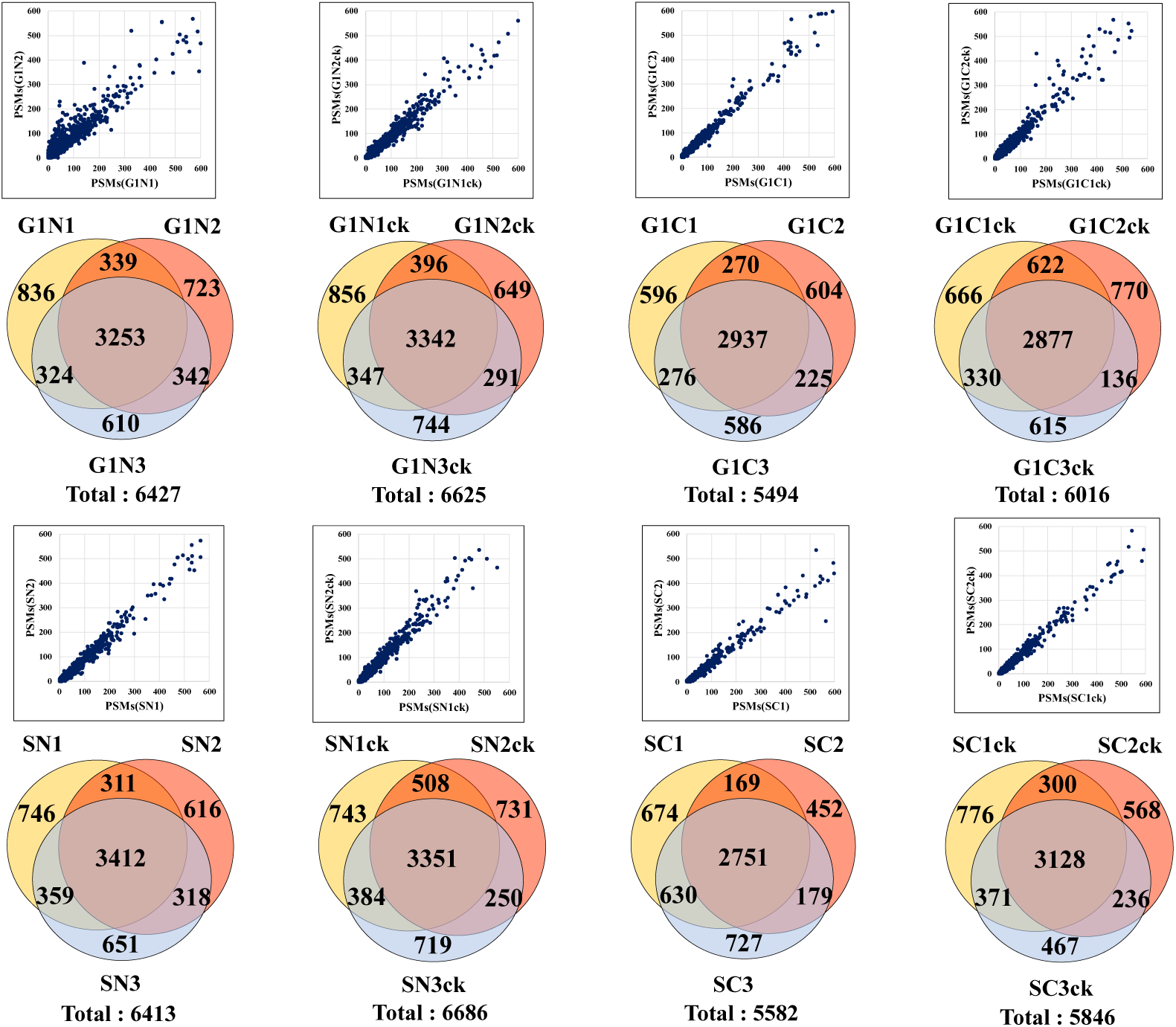
Reproducibility of protein identifications in microglia cells. Scatterplots show the correlation between the PSMs in two biological replicates, and the Venn diagrams show the protein overlaps in three biological replicates of G1, S and nuclear and cytoplasmic cell fractions.

The effectiveness of the nuclear protein enrichment step was assessed based on subcellular protein assignments provided by UniProt in the top 100 proteins with the largest number of distinct peptide counts (**Figure 3A**). On the average, the nuclear-enriched cell extracts contained ∼70 % nuclear proteins, vs. ∼33 % in the cytoplasmic fractions and whole cell/non-enriched lysates. The results were consistent throughout the entire dataset. Overlaps between the nuclear and cytoplasmic proteins were observed and expected to occur, due to the interference of the nucleocytoplasmic shuttling processes, linkage of the cytoskeletal to the nucleoskeletal proteins [51], and the association of various cytoplasmic components (transcription factors, signaling molecules, gene modulators, etc.) with the nuclear material. The reproducible detection of a specific set of nuclear and cytoplasmic “barcode” proteins that were shown previously to display a consistent number of spectral counts irrespective of the treatment performed on cells [52], as well as of a few proteins used as controls in routine biological experiments (e.g, tubulin, actin, GAPDH), further corroborated the quality of the data and reproducibility of the nuclear/cytoplasmic separation process **(Figures 3B** and **C)**. Classical nuclear and cytoplasmic markers were identified in proportion to their abundance in the expected cellular fraction **(Figures 3D** and **E)**.

**Figure 3.**
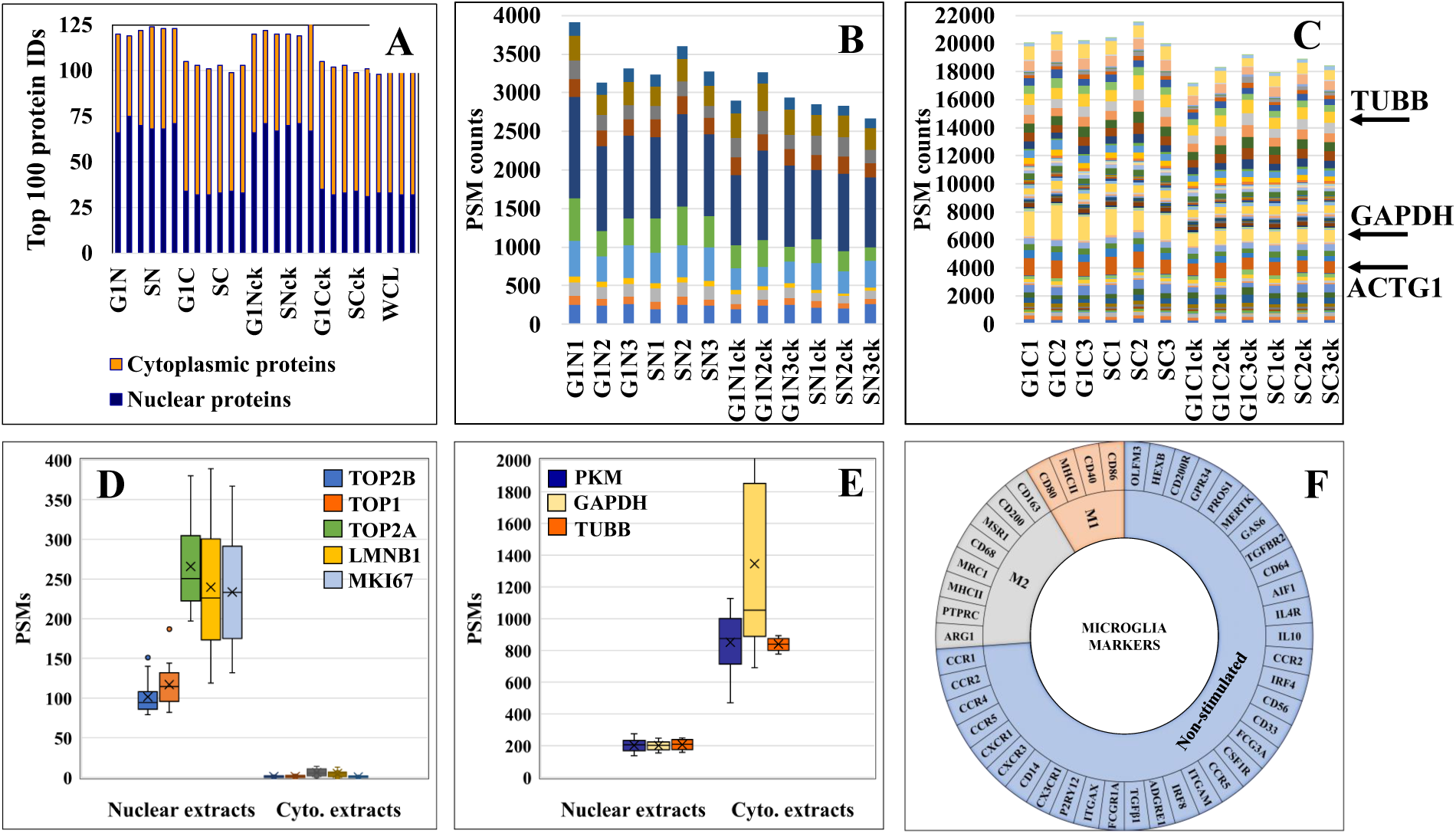
Charts representing the detection reproducibility of various protein marker categories in HMC3 cells. (**A**) Nuclear enrichment process outcome. The number of proteins with nuclear association in the top 100 most abundant proteins from the dataset is shown. Due to dual cellular component assignments, the sum of the nuclear and cytoplasmic proteins in the nuclear cell extracts exceeds 100. (**B**) Nuclear barcode proteins (11) displaying stable PSMs in the various nuclear enriched cell fractions [52]. (**C**) Cytoplasmic barcode proteins displaying stable PSMs in the various cytoplasmic cell fractions. Beta tubulin (TUBB), glyceraldehyde 3-phosphate dehydrogenase (GAPDH) and cytoplasmic actin gamma 1 (ACTG1) are classical cytoplasmic marker proteins. (**D**) Abundance of nuclear markers in the nuclear and cytoplasmic cell fractions (topoisomerases-TOP1, TOP2A and TOP2B; lamin B1-LMNB1; and proliferation marker protein-MKI67). (**E**) Abundance of cytoplasmic markers in the nuclear and cytoplasmic cell fractions (TUBB, GAPDH, pyruvate kinase isozymes M1/M2-PKM). (**F**) Sunburst chart of microglia marker proteins associated with immune-activation and cytokine signaling processes.

### Microglial markers in HMC3

A first priority in analyzing the HMC3 proteomic data revolved around assessing whether the cells presented protein signatures characteristic of microglia function and/or M2 polarization. Microglia have been extensively characterized for their cell surface markers to define their physiology and distinguish them from peripheral macrophages in the brain. Over 50 proteins have been catalogued to define different states of human microglia, most of which being associated with immune-activation and cytokine signaling processes [21,53] (**Figure 3F**). In this study, we identified 14 of these proteins, mainly associated with the plasma membrane, with known roles in regulating proliferation, differentiation, and immune functions in cells of the myeloid lineage **(Supplemental Table 3)**. One of the most relevant proteins indicative of M2 polarization was the macrophage scavenger receptor MSR1 (CD204) [4] that was identified by three unique peptides (7 PSMs) in the cytokine-treated cells, and by only one PSM in the non-treated cells. MSR1 is a cell membrane glycoprotein involved in host defense and microglia activation during neuroinflammation, and has scavenger activity in macrophages. Other markers such as the receptor-type tyrosine-protein phosphatase C (PTPRC or CD45) and arginase 1 (ARG1) [4] were identified in various cell fractions, but only by a single PSM. PTPRC has essential roles in regulating T- and B-cell antigen receptor and cytokine signaling processes, and ARG1 is a cytoplasmic enzyme of the urea cycle with roles in innate and adaptive immune responses and IL-4 signaling events [32]. Markers of M1 polarization such as CD 40, CD80, and CD86 [4,10,54], which are expressed upon treatment with pro-inflammatory cytokines, were either not detected or detected with very low or decreased spectral counts (e.g., MHCII/HLAII). Proteins that have been shown previously to be highly expressed or unique to microglia over other immune cells were consistently observed in both untreated and cytokine-treated cells [55,56]. These proteins included the purinergic receptor (P2RY12), anticoagulant plasma protein (PROS1), receptor tyrosine kinase (MERTK), olfactomedin-like protein 3 (OLFM3), beta-hexosaminidase subunit beta (HEXB), and the Fc receptor-like proteins (FCRLs). Moreover, proteins that were part of a top 10 adult microglia signature protein set, and that could not be identified in previously analyzed microglia cell lines [55], were identifiable in the human fetal brain-derived HMC3 cells [P2RY12, PROS1, MERTK, FCRLs, apolipoprotein E (APOE), sal-like protein 1 (SALL1), and integrin beta 5 (ITGB5)]. A few specific receptors, typically observable in non-stimulated or resting microglia cells, are worth additional comments. Remarkably, CD33, which belongs to the Siglec (sialic-acid-binding immunoglobulin-like lectin) family of molecules with roles in inhibiting immune cell activation and maintaining the cells in a resting state [57], was present only in the untreated HMC3/S cells but absent in the cytokine-treated cells. The enzyme-linked receptor quartet composed of MERTK, PTPRC, colony stimulating factor 1 receptor (CSF1R), and transforming growth factor beta receptor 2 (TGFBR2) were present in both treated and untreated datasets. These receptors have various regulating roles in cell proliferation, differentiation, survival, apoptosis, and inflammatory responses, and via MAPK, STAT and TGFβ signaling are also involved in oncogenic cell transformation. Other markers included three G-protein coupled receptors, i.e., adhesion G protein-coupled receptor E1 (ADGRE1) involved in cell adhesion and immune cell-cell interactions, and the C-C chemokine receptors type 4 and 5 (CCR4 and CCR5) with roles in chemokine signaling in macrophages. Also, the resting HMC3 cells were categorized by ATCC as negative for the GFAP (Glial fibrillary acidic protein) and positive for IBA1 (AIF1-Interferon gamma responsive transcript) markers. IBA1 was not detectable in the cells, but AIF1L (Allograft inflammatory factor 1-like), an AIF1 paralog, was upregulated in the serum-depleted/cytokine-treated cells. It was also interesting to note the substantial upregulation of GFAP in all cytokine treated cells. GFAP is a cytoskeletal protein, typically considered specific to astrocytes during the development of the CNS. Upon astrocyte activation, it was shown that GFAP expression is increased to inhibit inflammatory responses and reduce oxidative stress [58].

### HMC3 surfaceome and its relevance to immune functions

To gain an insight into the HMC3 cell surface proteome and its immunological implications, relevant categories such as receptors, transporters, and cell junction and adhesion proteins, along with other proteins with catalytic activity and cell surface antigens, were extracted from the dataset based on UniProt keyword and GO controlled vocabulary annotations (**Figure 4** and **Supplemental table 3**). From the total of 10,832 proteins, ∼8,000 were matched by two or more distinct peptides, and could be assigned to the general nuclear (2,895), cytoplasmic (3,026) and cell membrane (977) cellular sub-fractions. The HMC3 surfaceome, represented by 857 proteins mapped to specific cell membrane annotations, was captured in the dendogram shown in **Figure 4A**. This protein set represents ∼23 % of the UniProt annotated *Homo sapiens* cell membrane proteins (3,695), and could be assigned to all major categories, i.e., Tyr and Ser/Thr receptor and nonreceptor kinases, GPCRs, transporters, cell-cell and cell-matrix junctions, and to other matrix binding and interacting proteins comprised of GPI-anchored, matrix metalloproteases and scaffolding extracellular matrix glycoproteins. Together, these proteins are responsible for detecting extracellular cues and orchestrating the signal transduction process that triggers a corresponding cellular response, and represent the focus of drug target discovery efforts for therapeutic interventions.

**Figure 4.**
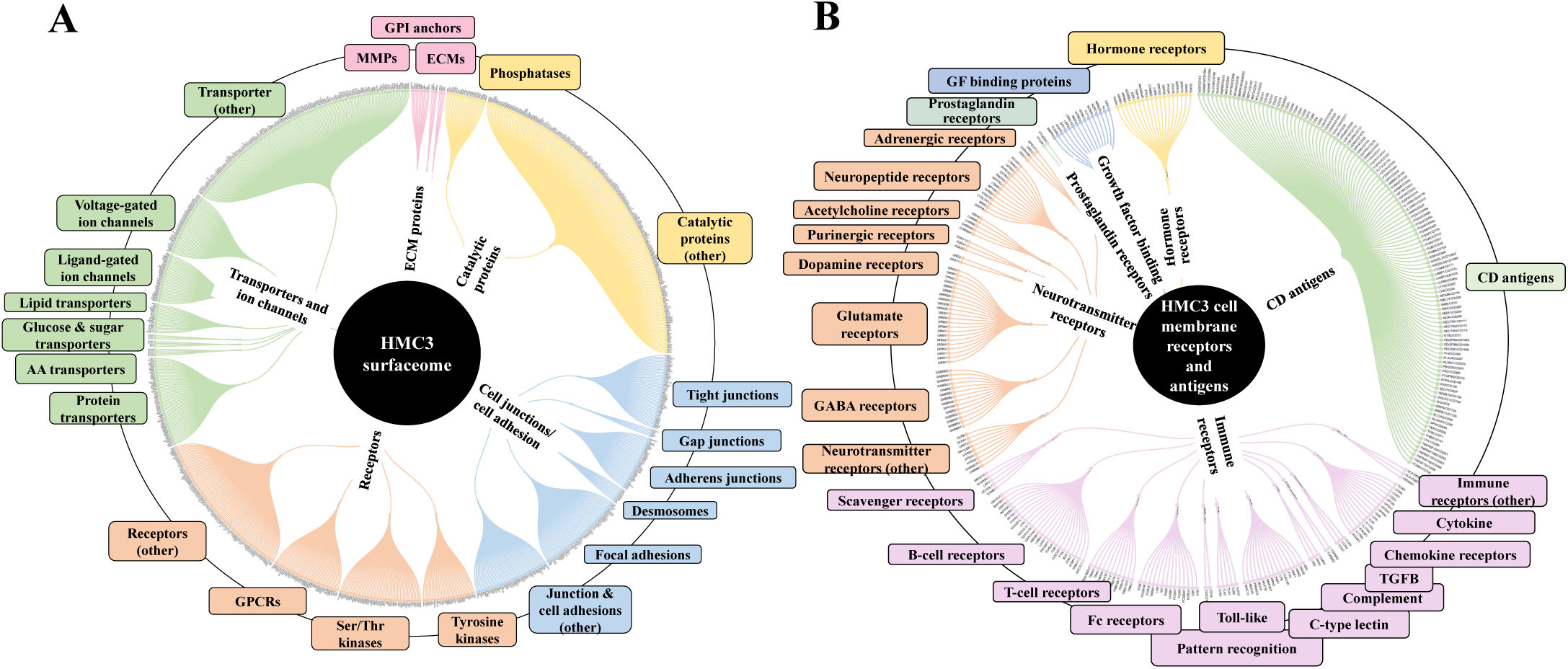
Dendograms representing the HMC3 microglia surfaceome. (**A**) HMC3 cell membrane proteins represented by receptors (Tyr and Ser/Thr receptor and nonreceptor kinases; olfactory, frizzled, taste, neurotransmitter and adhesion GPCRs), transport proteins (ligand and voltage-gated ion channels, transporters of small and large molecules), cell adhesion and cell-cell (tight junctions, gap junctions, adherens junctions and desmosomes) or cell-matrix (focal adhesion) junctions, and proteins with various catalytic functions (857 proteins with two distinct peptides, out of 1314 total cell membrane proteins detected; ∼18 % present in multiple categories). (**B**) HMC3 cell membrane receptors and CD antigens. A total of 283 proteins were assigned to signaling processes triggered by immune receptors (cytokine, chemokine, PRRs, complement receptors, Fc, T-cell, B-cell, and scavenger receptors), neurotransmitter receptors, hormone and prostaglandin receptors, glycan sensors (C-type lectin and Siglecs), and other growth factor binding proteins and receptors.

The entire landscape of specialized cell membrane receptors (growth factor binding, immune, neurotransmitter and hormone receptors) and CD antigens that initiate and carry out various microglial signaling functions was represented in the HMC3 cells. The different categories are highlighted in the dendrogram from **Figure 4B**, and the immune response processes that can be triggered by the stimulation of the HMC3 cells via these receptors, as well as the complex protein-protein interaction (PPI) networks between the receptors, in **Figure 5**. Infectious agents, pathogens, damaged/apoptotic or cancerous cells in the CNS release a variety of factors that trigger an inflammatory response from the host. Additional release of chemokines provokes receptor mediated signaling processes that promote the targeted movement of microglia toward the site of pathogenesis (chemotaxis), recognition of specific molecules on the surface of the target, and initiation of an immune response that eliminates the perturbation source, cellular debris and neurotoxic agents (phagocytosis). Chemotactic migration of microglia and solicitation of an immune response is the result of a complex interplay between purinergic receptors, ion channels, neurotranmitters and Tyr kinase signaling [59,60]. Neurotransmitter receptors (glutamate, GABA, acetylcholine) modulate a variety of microglia functions, including activation, phagocytic clearance and polarization [61]. GABA, acetylcholine and adrenergic receptors modulate and limit the release of inflammatory molecules and exert neuroprotective functions [60,61]. Stimulation of glutamate receptors has been shown to induce actin cytoskeleton rearrangements to promote chemotaxis and phagocytosis, and a separate group of the glutamate family of receptors modulate neurotoxicity or have neuroprotective functions [62]. Adrenergic receptors have been shown to control M2 activation by negatively affecting the expression of inflammatory IL-6, NO and TNFα in the cells [60,61,63]. Microglia use cell surface pattern recognition receptors (PRRs) such as toll-like receptors (TLRs) and C-type lectin receptors (CLRs) complexed with scavenger receptors (CD204, CD14, MERTK, AXL receptors) to detect and bind to specific pathogen-associated molecular patterns (PAMPs) on the surface of pathogens. Alternatively, microglia use TREM-2 receptors to recognize damage-associated molecular patterns (DAMPs) of apoptotic neural and other dead or damaged cells to initiate phagocytic clearance of the captured particles [64-66]. Phagocytosis is further regulated by Fc*γ*, complement, or purinergic receptors [67].

**Figure 5.**
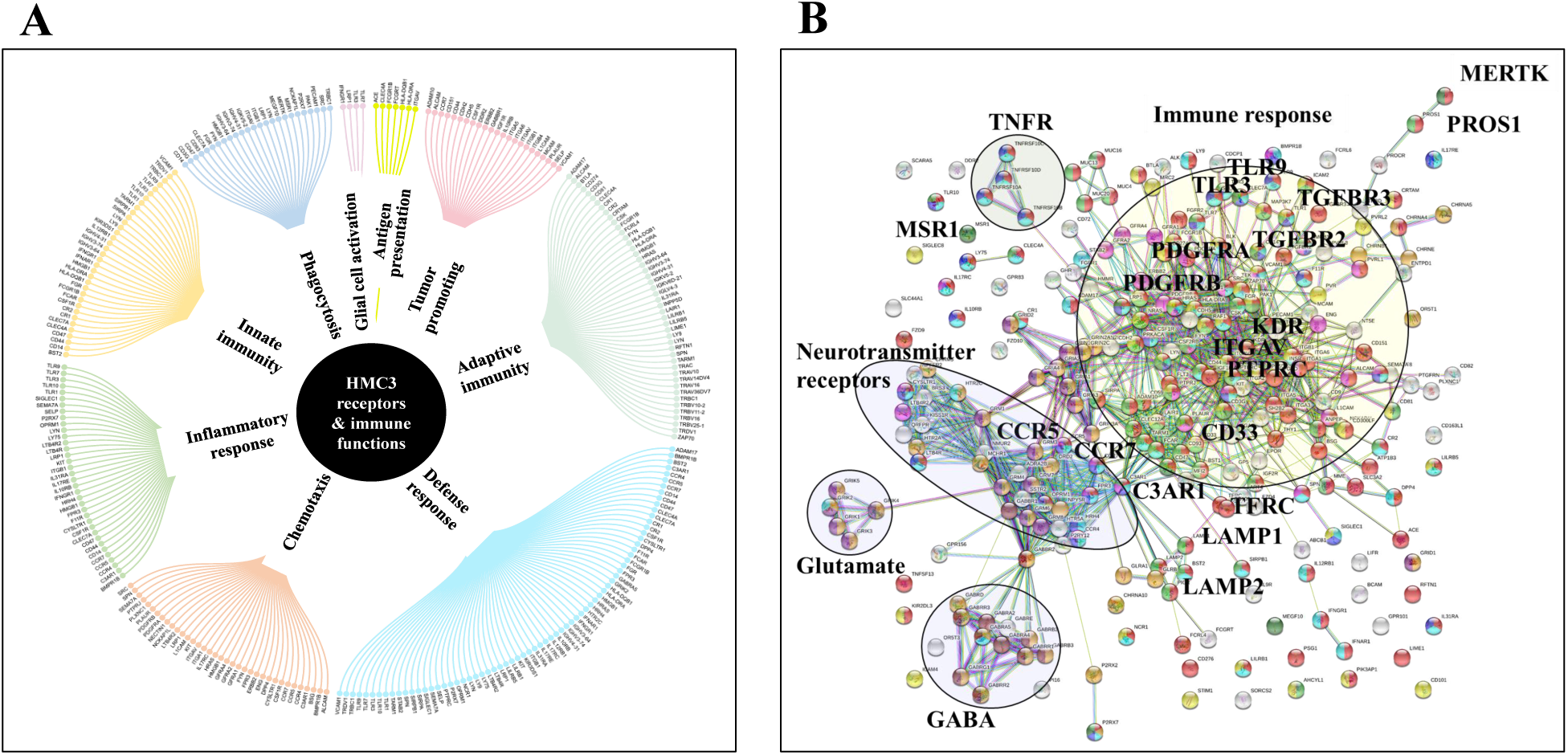
Immune response processes triggered by HMC3 cell membrane receptors and CD antigens. (**A**) Dendogram of major processes: innate and adaptive immunity responses, defense/inflammatory responses, chemotaxis, phagocytosis, secretion of immune modulators, antigen presentation, and tumor support. (**B**) STRING PPI diagram of the HMC3 receptors and CD antigens: red-immune system process, green-innate immune response; yellow-adaptive immune response, blue-inflammatory response, purple-chemotaxis; dark green-phagocytosis, light blue-defense response, dark yellow-chemical synaptic transmission, dark purple-glutamate receptor pathway, brown-gamma aminobutyric acid signaling pathway.

Downstream, specialized proteins such as ELMO1 (Engulfment and cell motility 1) and DOCK1 (Dedicator of cytokinesis 1) are activated to control cytoskeletal rearrangements that support phagocytosis and motility [68]. The internalization and degradation of the targeted entity is followed by presentation of the antigens to T-cells. The antigen processing and presentation cluster was comprised of a number of intracellular degradative enzymes such as cathepsins, ERAP, and legumain, along with proteasome subunits that are required by the cell for trimming peptides before presentation for binding to MHC molecules. Protein complexes that transport vesicles containing digested antigens along the microtubules and localize them at the cell membrane were observed in high numbers. MHC/HLA class I molecules that enable antigen presentation to the adaptive cytotoxic T-cells were detectable [69]. Altogether, the interplay between the various receptor systems bridge innate and adaptive immune responses implicated in defense, inflammation and cytokine production, via the activation of various signaling pathways such as TLR, MAPK, ERK1/2, PI3K/AKT, JNK, JAK/STAT, and NFKB. These capabilities were all supported by the receptors identified in the HMC3 surfaceome.

### Comparative assessment of differentially expressed proteins

Comparative proteomic analysis was performed between cytokine-treated and non-treated cells, in the G1 and S cell states. The Volcano plots from **Figure 6** depict the respective comparisons, i.e., G1Nck vs G1N, G1Cck vs G1C, SNck vs SN, and SCck vs SC, with up/downregulated biological processes encompassing protein counts of 433/307, 332/113, 484/374 and 260/102, respectively. Some cell membrane proteins were detected in the nuclear fractions due to co-precipitation during centrifugation. Only proteins identified with a minimum of two unique peptides were considered for downstream bioinformatics analysis. The biologically relevant differences arising due to the treatment of cells with anti-inflammatory cytokines were explored using STRING, DAVID, KEGG, Reactome, UniProt and GeneCards bioinformatics identification, annotation and enrichment tools (**Figures 7** and **8**). **Supplemental Tables 4** and **5** include the full lists of proteins, enriched protein categories, and associated networks and pathways. For a cohesive biological interpretation, enriched GO categories from the nuclear and cytoplasmic fractions of a cell state were combined and discussed collectively. In addition, categories of relevance to immune-response were explored individually (**Figure 7**). Up- and downregulated proteins and biological processes were observed in all cell states (**Figure 7A**), with observable impact from cell-surface-induced to transcription factor activities (**Figure 7B**), and output affecting cellular communication/activation, transcription/translation, transport, metabolic processes, cell cycle, growth, cell death (**Figure 7C**), and various immune responses (**Figure 7D**). General categories related to nucleic acid, protein or small molecule metabolism, as well as transport and signaling, encompassed large protein groups that were observable in all comparisons that were explored. As regulation of protein activity and function is often achieved at the post-translational level, the addition or removal of PTMs alters the detectability of proteins. This can be especially the case of proteins involved in signal transduction and cell cycle regulation, for which the observed changes may have been representative of changes not just in expression level but also in activity. Specific categories that are discussed below were selected based on relevance to the cytokine treatment that was performed.

**Figure 6:**
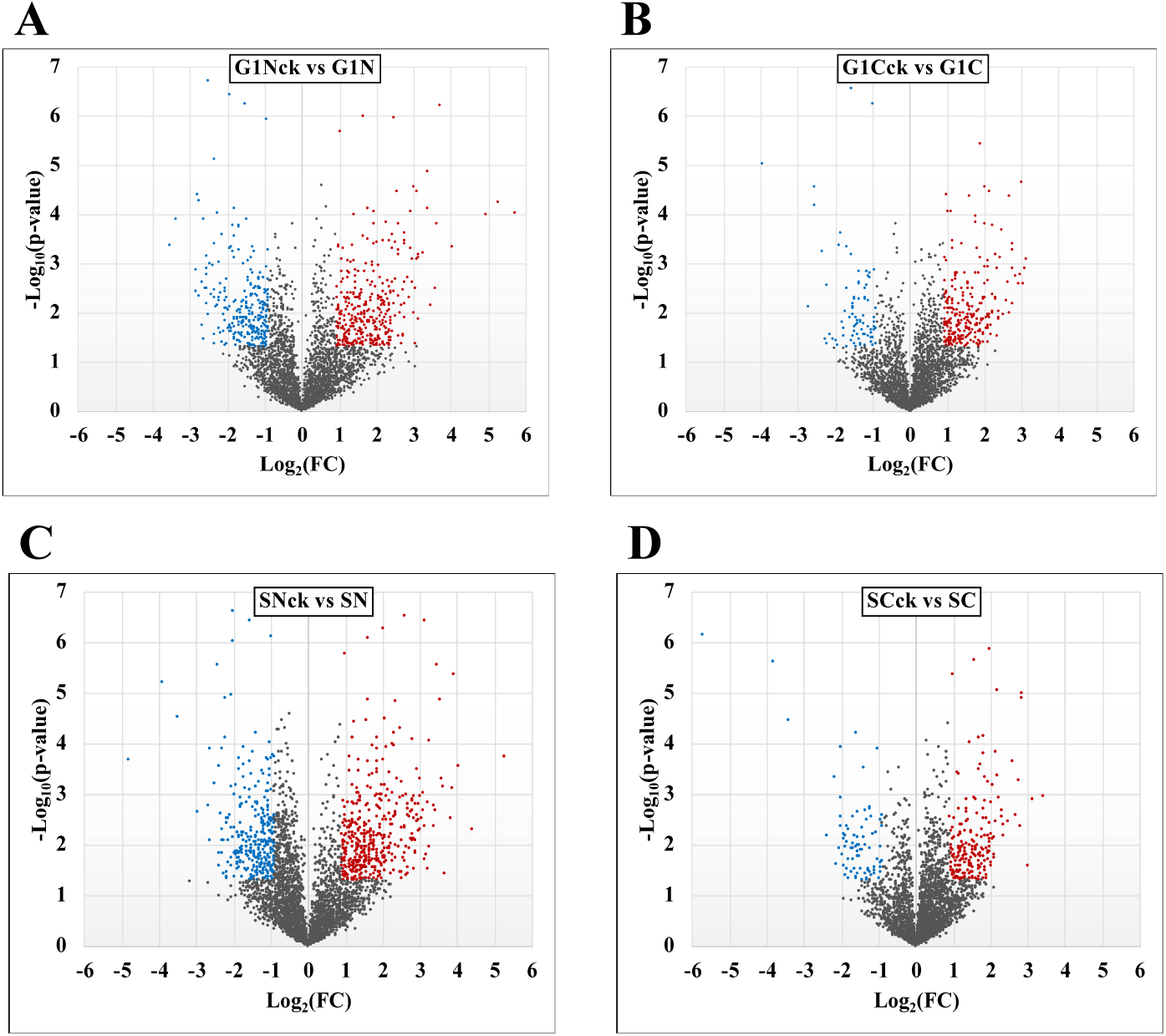
Volcano diagrams depicting proteins associated with differentially modulated pathways in cytokine-treated vs non-treated HMC3 cells.

**Figure 7:**
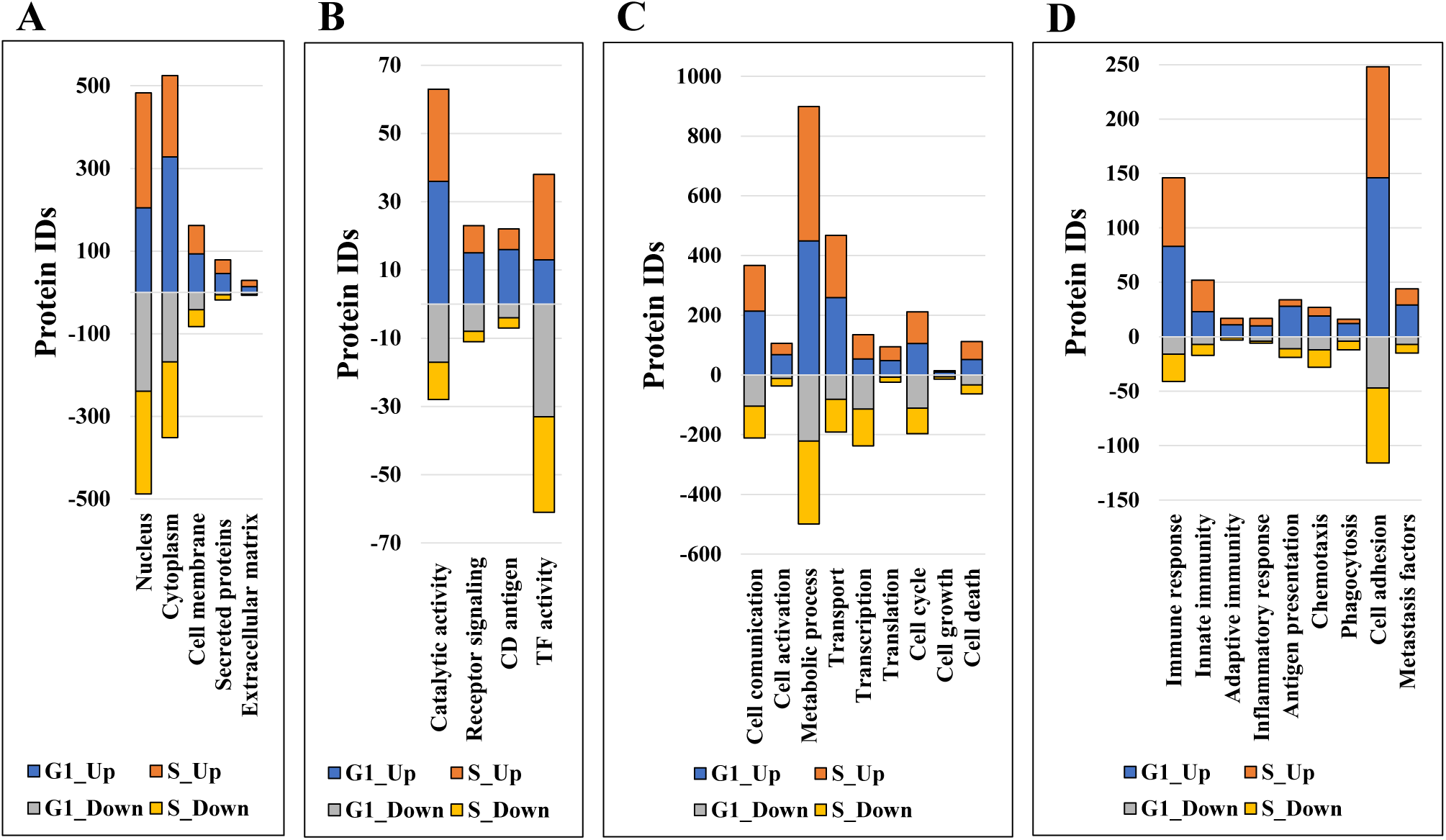
Stacked bar charts representing proteins that changed expression or activity level and could be associated with general biological categories related to: (**A**) Cellular location, (**B**) Receptor/catalytic or TF activity, (**C**) Biological processes, and (**D**) Immune response.

## Discussion

### Upregulated biological processes in serum-depleted/cytokine-treated cells

The upregulated processes in the serum-depleted/cytokine-treated cells were represented by a total of 728 proteins clustered into four broad biological categories, i.e., (a) mitochondrial gene expression and oxidation-reduction, (b) ECM remodeling and cell adhesion/migration, (c) immune response, and (d) regulation of cell cycle. **Figure 8A** provides a representative selection of these biological processes, **Figure 9A** a STRING PPI network of the whole protein set, and **Figure 10** a breakdown of the PPI network into the above four categories.

**Figure 8:**
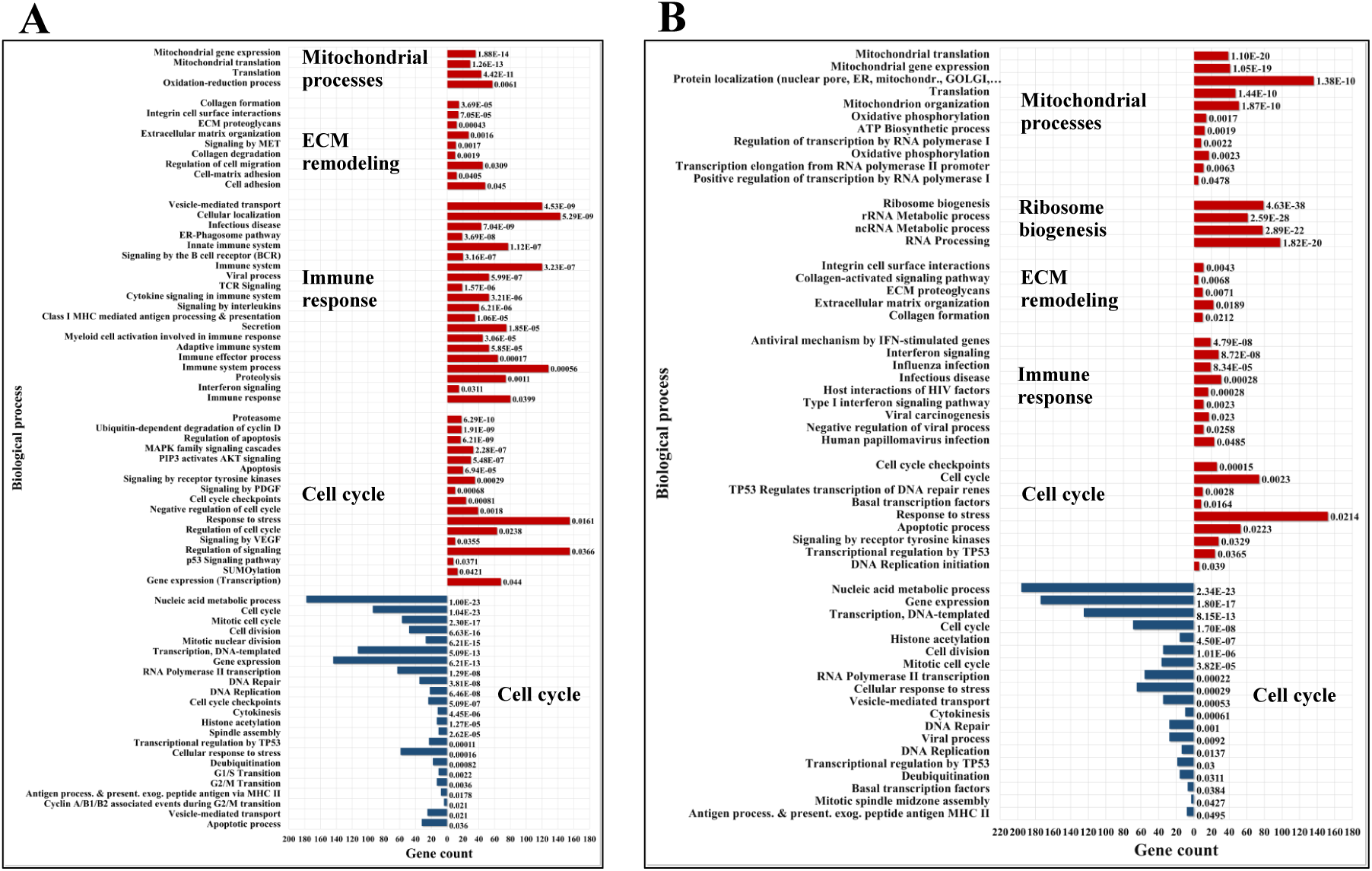
Bar charts of selected enriched up-(red) and down-(blue) regulated biological processes and pathways in cytokine-treated vs non-treated HMC3 cells. (**A**) G1ck vs G. (**B**) Sck vs S. Annotations are based on BP, KEGG and Reactome databases. Full lists of enriched categories are provided in **Supplemental Table 5**.

**Figure 9.**
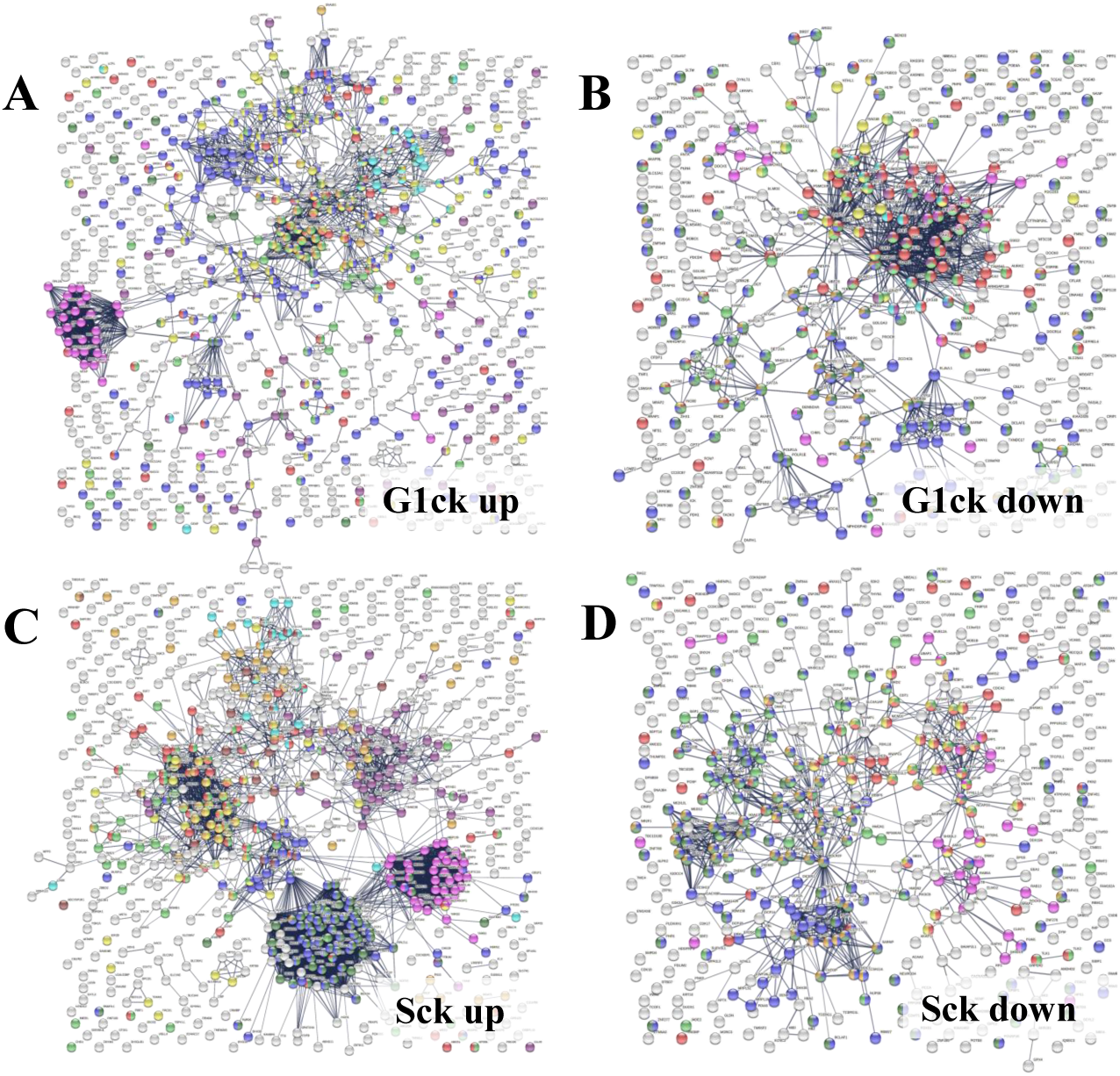
STRING PPI networks representing up- and downregulated biological processes in cytokine-treated vs. non-treated HMC3 cells. **(A) Upregulated G1ck vs. G1 (728)/Details provided in Figure 10:** Magenta-Mitochondrial gene expression (translation) (36); Purple-Oxidation reduction process (60); Yellow-Immune system process (127); Red-Regulation of cell cycle (63); Green-Proteolysis/phagosome pathway/FCERI mediated NFKB activation (74); Turquoise-ECM organization (27); Blue-Transport/vesicle mediated transport (228); Dark yellow-MAPK family signaling (33); Dark green-Regulation of cell migration (44). **(B) Downregulated G1ck vs G1 (403)**: Red-Cell cycle (94); Green-Chromosome organization (82); Blue-Gene expression/transcription (144); Yellow-DNA repair (35); Dark green-Transcription DNA templated (113); Dark yellow-Transcription RNA pol II (40); Purple-M phase (23); Turquoise-S phase (13); Magenta-Membrane trafficking (24). **(C) Upregulated Sck vs S (714)**: Magenta-Mitochondrial gene expression/translation (mitochondrial ribosomes) (41); Purple-Mitochondrial organization/oxphos, ATP biosynthesis, respiratory electron transport (51); Dark green-Ribosome biogenesis (79); Blue-RNA processing, rRNA/ncRNA (98); Yellow-Viral processes/Interferon signaling (51); Red-Cell cycle (74); Green-Chromosome organization (67); Turquoise-ECM organization (22); Dark yellow-PTMs (69); Brown-Signaling by RTK/cytoskeletal rearrangements, membrane organization, integrin cell surface interactions (28). **(D) Downregulated Sck vs S (465)**: Red-Cell cycle (69); Green-Chromosome organization (77); Blue-Gene expression/transcription (174); Yellow-Mitotic cell cycle process (36); Dark green-Transcription DNA templated (125); Dark yellow-RNA polymerase transcription II; (56) Magenta-Membrane trafficking (35).

**Figure 10.**
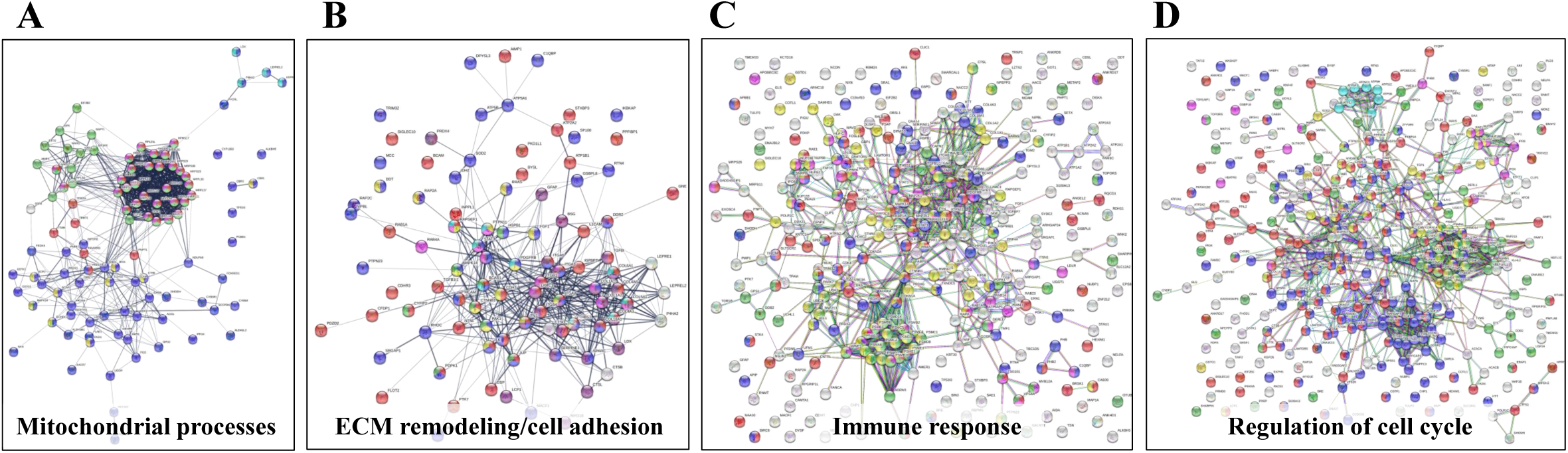
Detailed STRING PPI networks representing the upregulated biological processes in serum-depleted/cytokine-treated HMC3 cells (G1ck vs G1). The networks were built based on the selected enriched protein categories shown in Figure 8. (**A**) **Mitochondrial gene expression and redox processes**: Red-Mitochondrial gene expression (36); Blue-Oxidation-reduction process (57); Green-Translation (42); Magenta-Mitochondrial translation (30); Yellow-Fatty acid metabolic process; (9) Turquoise-Collagen formation (4). (**B**) **ECM remodeling and cell adhesion**: Red-Cell adhesion (48); Blue-Regulation of cell migration (45); Purple-Extracellular matrix organization (27); Yellow-Regulation of MAPK cascade (15); Magenta-Signaling by Met (11); Turquoise-Signaling by PDGF (9); Green-Signaling by VEGF (9). (**C**) **Immune response**: Red-Immune system process (128); Blue-Vesicle mediated transport (120); Green-Proteolysis (74); Magenta-Protein localization to the nucleus (19); Turquoise-Mitochondrial ATP synthesis coupled proton transport (8); Yellow-Cytokine signaling in immune system (53); White-Localization. (**D**) **Regulation of cell cycle**: Red-Regulation of cell cycle (63); Blue-Regulation of apoptotic processes (73); Green-Proteolysis (55); Yellow-Immune system (83); Magenta-Viral processes (39); White-response to stress/signaling processes/localization. **Selected proteins associated with the observed biological processes. mt-RNA processing/translation**: TFAM, TFB2M, TRNT1, AARS2, GFM1, TUFM, PNPT1, KIAA0391; **FAO**: HSD17B10, ACAA1, ECHS1, ECI1, ETFB; **Mitochondrial and cytoplasmic oxidoreductases**: PRDX3/4/5/6, GSTK1, GSTO1, SOD2; **Electron transport chain**: NADH dehydrogenases NDUFA8, NDUFAF3, FOXRED1; **Succinate dehydrogenase** SDHA; **ATPase subunits**: ATP5D, ATP5F1A, ATP5MF, ATP5PO, ATP5PF, ATP5F1C, ATP5F1B, ATP5PB); **Collagens**: COL1A1/2, COL4A3/6, COL5A1, COL6A1, COL18A1; **MAPK signaling**: MAPK14, PAK1, PTPN11, ILK, FGF1, CTNNB1, RRAS, RAP2A, TAB1; **MET signaling**: MET, PTPN11, LAMC1, RAB4A, STAT3; **PDGF signaling**: PDGFRB, RAPGEF1, BCAR1, PTPN11, STAT3; **VEGF signaling**: PAK1, CTNNB1, JUP, MAPK14; **TGFβ signaling**: TAB1, ZFYVE9, PAK1; **Integrin signaling**: ITGA3, ITGA6, ITGAV, ITGB1, L1CAM, MET; **Cell surface proteins and receptors**: PDGFRB, BSG1, DDR2, PTK7, ITGAV, ITGB1, CDHR3, TMEM33; **Intracellular signaling effectors**: BCAR1, JUP, PALLD, TGFB1I1, FYN, RRAS, RAPGEF1, RAP2C, ARHGAP24, ARHGEF18, CTNNB1; **Cell adhesion and junction proteins**: MCAM, L1CAM, BCAM, CDH2, CDHR3, THBS1, TGFBI, IGFBP7, CTNNB1, VCL, JUP, DSP; **ECM remodeling enzymes**: CTSB, CTSD, CTSL, LOX, NEU1, FUCA1; **Other ECM molecules**: SPARC, SERPINE1, FN1, TNC, LAMC1, BSG; **Cytokine signaling**: MYD88, TAB1, PAK1, PTPN11, CDKN1A, PML, FYN, nuclear import, proteasome components and ubiquitin ligases); **Aminopeptidases**: LNPEP, NPEPPS, ERAP1/2; **Golgi vesicle transport**: TMED2/7/9/10, COPI coat (COPB1/2, COPE, COPG1), AP-1/AP-4 (AP1G1, AP1S2, AP4M1) complexes; **Vesicle fusion to the target membranes**: ARFGAP1/3, EXOC4, SNARE complex (NAPA, STX12, YKT6); **Ubiquitin-protein E3 ligases**: UBE3A, UFL1, RNF213, WWP1, KLHL2, TRIM25 and TRIM32; **Cell cycle arrest/negative regulation of growth**: CDKN1A, CDKN2A, CEPBA, GADD45GIP1, CDK4, CDK6, PML, ILK, TSG101, PHB. **Downregulated cell cycle processes. Spindle assembly**: AURKB, AURKC, INO80, KIF2A/4A/4B/C1, NEK6, RACGAP1, TPX2, TUBGCP6; **DNA repair and replication**: BRCA1/2, ERCC1, LIG3, RAD51, BARD1, BLM, CHAF1A, MCM10, MEN1, POLE, POLD2, POLK, RFC3/5, ORC3/4, TOPBP1.

#### Mitochondrial gene expression and oxidation-reduction

Cells produce energy by oxidizing their carbon fuels in the TCA cycle and generate a proton-motive force by repeated oxidation-reduction cycles to generate ATP in a process known as oxidative phosphorylation (OXPHOS). This process occurs in the mitochondria. Noticeably, the cytokine-treated HMC3 cells showed an upregulation of mitochondrial gene expression reflected by proteins involved in mitochondrial transcription, mt-RNA processing and translation, and an increased expression of 25 mitochondrial ribosomal (MRP) protein subunits including 18 large and 7 small (**Figure 10A**). These results were complementary to the identification of various protein complex subunits implicated in mitochondrial and cytoplasmic oxidation-reduction reactions, oxidative phosphorylation, cellular response to oxidative stress, cell detoxification, and lipid/organophosphate/organonitrogen compound metabolism. It is now well established that IL-4 treated M2 microglia are characterized by lower glucose consumption by switching to oxidative phosphorylation which is fueled by higher rates of fatty acid oxidation (FAO) [70]. Accordingly, an upregulation of mitochondrial enzymes that are involved in FAO was observed. In addition, the reactive oxygen species (ROS) generated during oxidative phosphorylation were balanced by an upregulation of mitochondrial and cytoplasmic oxidoreductases which protect the cells during oxidative stress.

#### ECM remodeling and cell adhesion/migration

Increased mitochondrial respiration in response to IL-4 is correlated with switching the arginine metabolism from citrulline and nitric oxide production in M1 microglia to arginase-catalyzed production of ornithine and polyamines for collagen synthesis in M2 microglia [38,41,71] (**Figure 10B**). Subsequently, the prolyl-hydoxylase enzymes (P3H1, P3H3, P4HA2) which are key to collagen synthesis showed an increased expression in the cytokine treated cells. Indeed, HMC3 showed a substantial upregulation of collagens, collagen receptors (ITGB1/ITGA3, DDR2), and cell-collagen adhesion modulators such as the TGFβ induced protein (TGFBI). Collagen deposition and cell-collagen interactions has been linked to cell invasiveness, and in cancer cells, to enhanced metastatic abilities [72]. The integrin ITGB1/ITGA3 heterodimers are receptors not just for collagen, but also for fibronectin, laminin and thrombospondin, and along with the ITGB4/ITGA6 receptors for laminin mediate invasiveness [32]. TGFBI, that binds to ECM and integrins, on the other hand, has been shown to inhibit cell adhesion [32]. Likely the result of co-stimulating these cells with TGFβ [38], MAPK signaling and regulation which is central to numerous cellular pathways, was noticeably altered in the cells. Along with MAPK, signaling by c-MET, PDGF, and VEGF were tightly interconnected to the upregulated processes related to ECM remodeling, collagen formation, cell-cell/cell-matrix adhesion, and cell migration. Together, the cell adhesion molecules, the cell surface proteins and receptors, and the cytoskeletal proteins and intracellular signaling effectors orchestrate cytoskeleton organization and remodeling, leading to enhanced cell migration. A deeper look at the extracellular enzymes (LOX, CTSB, CTSD), growth factors (FGF1, NGF), cytokines (AIMP1), and ECM molecules detected in the cells, highlighted their role in the extracellular matrix dynamics. ECM reorganization, increased deposition of collagen and glycoproteins such as fibronectin (FN1) and tenascin (TNC), which were substantially upregulated in the treated cells, result in increased rigidity and tensile strength of tissue and favor tumor growth, progression and invasion [73,74]. Enzymes such as cathepsins (CTSB, CTSD), SPARC (Secreted protein acidic and rich in cysteine), and protease inhibitors such as SERPINE1 (Serpin family E member 1), support tumor invasion and metastasis by interacting with, degrading and/or re-shaping the extracellular matrix [73]. LOX (Lysyl oxidase) expression, responsible for the oxidative deamination of Lys residues in collagen precursors, collagen crosslinking and increased matrix stiffening, was correlated with increased tumor adhesion and migration [73,76], but also with possible roles in tumor suppression [32]. Proteases (CTSB, CTSD) that participate in protein degradation and turnover activate tumorigenic invasion and metastatic pathways by cleaving signaling peptides, growth factors and hormones in the intra- and extracellular environment [77]. β-catenin (CTNNB1 or Cadherin associated protein, Beta 1), an important downstream effector of canonical Wnt signaling, was found to be consistently upregulated in all cytokine treated fractions, but mostly in the nuclear ones, pointing toward Wnt signaling activation. It has been well documented that TGFβ reciprocally induces the activation of Wnt ligands [78], and SMAD proteins of the TGFβ pathway engage in a crosstalk with the Wnt signaling proteins to release CTNNB1 from the GSK3β mediated destruction complex, stabilize its function, and regulate gene expression [79]. Recent studies have also shown that Wnt ligands produced by cancer cells use β-catenin signaling to stimulate immunosuppressive TAMs that support tumor growth and metastasis, and to upregulate the production of anti-inflammatory IL-4 and TGFβ [80]. However, both positive and negative regulators of the Wnt pathway, such as AMOTL1 (Angiomotin like 1), AMER1 (APC membrane recruitment protein 1-a positive and negative regulator of Wnt pathway), and LZTS2 (Leucine zipper tumor suppressor 2), were present alongside β-catenin, advocating for a complex method of β-catenin activation in these cells. IL-4 stimulation of macrophages, for example, was shown to induce the expression of E-cadherin which is stabilized by β-catenin to regulate cell adhesion processes [81]. E-cadherin was not detected in the cells, but the cytoplasmic fractions were highly enriched in the adherence junction cadherin modulators CDH2, CDH4 and CDHR3 (a viral receptor). Increased expression of cadherins boost homophilic cell-cell adhesion between cells, impeding therefore single cell migration, but facilitating grouped migration and cell motility on cadherin substrates. Cadherin molecules can also activate intracellular signaling pathways and alter the actin cytoskeleton via the various catenins (α, β and p120) that they bind [82]. In addition, the overexpression of focal adhesion proteins such as integrins, their regulator ILK (Integrin-linked kinase), talin modulators of integrin activation (TLN1/TLN2), and vinculin (VCL), indicated that integrin mediated signaling was highly correlated with the ECM processes and involved in transducing the extracellular cues into biochemical intracellular signals that alter cytoskeleton organization and migratory capabilities. Integrin molecules such as ITGAV serve as receptors for vitronectin (VTN), tenascin-C (TNC), thrombospondin (THBS1), fibronectin (FN1), fibrillin and laminin proteins, and together augment cell adhesion and signaling.

Enriched protein clusters that regulate chemotaxis or phagocytosis (e.g., TLR, P2X, P2Y, Fc or CCRX receptors) were either not identified, or identified with very low counts. Nevertheless, several receptors, cell adhesion and migration proteins in the stimulated cells could be associated with chemotactic processes (PDGFRB, VGFR1, DPYSL2, CRMP1, MET, PALLD, ITGAV, BCAR1, FYN, L1CAM). In addition, the detection of AIF1L (albeit with low counts), a paralog of AIF1 with roles in inflammation, macrophage activation, migration and induction of phagocytosis [31], pointed to the presence of such propensities. Overall, the combined effect of the upregulated proteins clearly demonstrated the induction of a microglia phenotype with enhanced ECM-remodeling and migratory capabilities, that simultaneously also directs the formation of a tumor-supportive ECM niche in the vicinity of the microglia cells [73,83].

#### Immune response

Proteins representative of immune reactions could be associated with innate, adaptive or inflammatory responses via cytokine signaling, class I MHC mediated antigen processing and presentation, chemotaxis, cell adhesion/migration, and vesicle mediated transport (**Figure 10C**). One of the mechanisms by which the type I and type II cytokine receptors signal is via the JAK/STAT pathway. The signal transducers and transcription activators STAT1 and STAT3 are known to get activated by interleukins and interferons [84,85], and complementary and conflicting roles of STAT1 and STAT3 on cell cycle have been also documented. STAT1 negatively regulates cell proliferation, elicits anti-viral and anti-tumorigenic immune responses, and promotes apoptosis. STAT3 promotes cell survival, G1/S transition and proliferation, motility and immune tolerance [32,86]. It has been shown, however, that increased expression in STATs does not necessarily correlate with increased activity [87]. JAK/STAT signaling pathways play critical roles in both innate and adaptive responses, with IL-4/IL-13 and IL-10 being known for downregulating NFKB and STAT1 activities, limiting the production of inflammatory cytokines, and driving an anti-inflammatory macrophage phenotype [39,88]. In response to IL-10 stimulation, STAT3 alone can mediate various cellular responses and inhibit the transcription and expression of pro-inflammatory cytokines [89], however, previous reports have suggested that in certain circumstances IL-10 can also activate STAT1 mediated signaling to instigate an inflammatory response [87]. These trends were observable in the present dataset, as well, and manifested themselves by the induction of M2a (via IL-4/IL-13) and M2c (via IL-10) polarized populations of cells with tumor-promoting activities that encompassed type II inflammatory/Th2 responses in M2a cells, and, matrix deposition, ECM remodeling and suppression of immune response in M2c cells [39]. STAT1 was found to be consistently upregulated in all cytokine-treated cells, but mostly in the G1Nck cell fractions, while changes in the activity of STAT3 were observable only in the G1Nck fractions, and appeared to be rather affected by the dephosphorylation of some of its matching peptides, including one that induces transcription upon phosphorylation at Y45 [90]. Nonetheless, a protein that mediates STAT3 degradation was also present (TMF1). STAT6 that is involved in initiating cytokine-initiated STAT signaling, was detectable, but with too low counts to support conclusive assessments.

Along with JAK/STAT, additional proteins participating in cytokine, integrin, MAPK, and TGFβ signaling (TAB1, ZFYVE9, SMAD) were highly interconnected in orchestrating the same cytoskeleton re-organization, ECM remodeling, adhesion, migration and endocytic processes. Binding of integrins to talin, vinculin, ILK, α-actinin, and filamin established the linkage between focal adhesions and the cytoskeleton to initiate integrin-mediated cell-ECM adhesions and in-and-out cell signaling. One of the downstream signaling cascades involved the docking protein BCAR1 (Breast cancer anti-estrogen resistance 1 scaffold protein or p130Cas). BCAR1 is an adaptor protein with roles in coordinating cell adhesion and inducing cell migration due to its activity in Tyr kinase signaling processes enabled by its multiple PPI domains and Ser/Tyr p-sites that bind SH2/SH3 domains of signaling proteins. Its unphosphorylated form localizes in the cytoplasm, the cell fraction in which it was observed with increased counts in this dataset [32]. TGFβ activated kinase 1 (TAB1) emerged as a signaling intermediate between TGFβ, Wnt and MAPK signaling, and PAK1 (P21-activated kinase 1) as an activator of downstream MAPKs. Activated by upstream integrin receptors, PAK1 plays roles in cytoskeletal reorganization, focal adhesion, migration, vesicle-mediated transport and chemotaxis. TGFβ signaling was further represented by the early endosomal zinc finger motif containing protein ZFYVE9 that recruits SMAD2/SMAD3 to the TGFβ receptor and mediates the subcellular localization and transcriptional activity of SMADs. SMAD2/4 were detectable in the data set, but with decreased counts in the nuclear fraction of cells. SMAD4 facilitates the binding of SMAD protein complexes to DNA to induce transcription, its regulation being heavily dependent on phosphorylation [32]. Phosphorylated peptides from complex cell extracts are difficult to detect by MS without prior enrichment because the addition of a phosphate group to the peptide backbone reduces the signal intensity substantially. In the present context, SMAD4 phosphorylation would have resulted in decreased counts, being indicative of TGFβ signaling activation. The concomitant downregulation of SMAD nuclear interacting protein SNIP1, who’s over expression has been shown to inhibit the formation of SMAD4 complexes and TGFβ signaling, reinforced this hypothesis. MYD88 (Myeloid differentiation primary response protein), an innate immune signal transduction adaptor involved in toll-like receptor signaling, was identified by few spectral counts, but changes in these counts were suggestive of altered NFKB activation. Changes in the counts of LCP1 (Lymphocyte cytosolic protein 1), an actin binding protein with roles in NFKB signaling and triggering T-cell responses [32], were also observed, but an impact on the expression of cell surface antigens that it modulates, IL2RA (Interleukin-2 receptor subunit alpha) and CD69 (Early activation antigen), could not be confirmed as these antigens were not detectable. Overall, NFKB signaling did not emerge as being activated in the serum-depleted/cytokine treated cells.

Class I MHC mediated antigen processing and presentation of ubiquitin-proteasome degraded proteins was represented by the HLA class I histocompatibility antigens (HLA-C Cw-8α and Cw-14α chains), various aminopeptidases that trim HLA class I-binding precursors for presentation by MHC class I molecules, and proteins with roles in folding (TCP1, CCT2, CANX), ER associated degradation (DERL1), activation of innate response pathways (MYD88), and degradation of proteins in the lysosomes (cysteine proteinase CTSL). Proteins belonging to the ubiquitin-mediated proteasomal degradation pathway that control a variety of immune functions including generation and processing of peptides for antigen presentation on MHC I molecules [91], were also observable. These included various subunits of the proteasome along with ubiquitin conjugating enzymes UBE2K and UBE2Z, and ubiquitin-protein E3 ligases. The proteasome components were amply overexpressed, and part not just of the antigen processing cluster but essentially of all global innate/adaptive response and signaling processes pertaining to cytokine, Fc epsilon receptor, CLR, MAPK and NFKB pathways. Interesting to note was the upregulation of TRIM25 which is an E3 ubiquitin/ISG15 ligase, where ISG15 is an interferon induced ubiquitin-like protein that activates or deactivates various targets involved in viral responses, and that can also induce the secretion of IFNG and IL-10 and act as a chemotactic factor for neutrophils [32]. The proteasome hub was equally prominent in the cell cycle cluster, and of relevance was the finding that components of the immunoproteasome (i-proteasome) that have essential functions in processing class I MHC peptides were present in both control and treated cells along with subunits of the constitutive 20S core [92]. The i-proteasome low molecular weight proteins LMP2 (PSMB9), LMP7 (PSMB8), and MECL-1 (PSMB10) were present concurrently with their standard proteasome subunit counterparts (i.e., PSMB6, PSMB5 and PSMB7) and the proteasome activator complex subunits 1, 2 and 3 of the 11S regulator of the i-proteasome (PSME1, PSM2 and PSME3). Immunoproteasome components can be induced by pro-inflammatory cytokines, but in the case of HMC3 cells appeared to be expressed at the basal level.

The immune cluster included a major overarching hub that comprised vesicular transport, viral and cellular localization proteins. Vesicular mediated transport plays a key role in a variety of immune related functions such as endocytosis of apoptotic cells and pathogens, exocytosis, cytokine secretion and antigen presentation. Main drivers of intracellular membrane trafficking, exocytosis and secretion (small Rab GTPases, GTPase activating ARFGAP1/3, EXOC4), Golgi vesicle transport (COPI coat and AP-4 complexes), fusion of vesicles to the target membranes (NAPA, STX12, YKT6), and transmembrane trafficking (TMED2/7/9/10), were all markedly overexpressed. Viral transport was mediated through integrin, proteasome and nuclear import (nucleoporins) processes. Several proteins such as GBP1 (Guanylate binding protein 1), PML (Promyelocytic leukemia protein/nuclear body scaffold), SP100 (SP100 Nuclear antigen), and MAVS (Mitochondrial antiviral signaling protein), were present to mount defense responses. Representative proteins of receptor-mediated endocytosis included DNAJC13, CANX, TFRC, LDLR and SPARC.

As the observed chemokines were detectable by only one peptide and one or two spectral counts, changes in chemokine expression could not be detected and measured with confidence. Nevertheless, chemokines CCL22, CCL23, CXCL12, and CXCL6 were observable only in the cytokine treated cells, and CXCL1 and CCL25 in both control and treatment. CCR4, the chemokine receptor for CCL22, was also detected only in the cytokine treated cells, while receptors CCR5 and CCR7 only in the control. With roles in immune surveillance and inflammation response, CCL22 is a chemoattractant for activated T-lymphocytes, NK cells, monocytes and dendritic cells, CXCL12 (SDF1) for T-lymphocytes and monocytes, and CXCL1 and CXCL6 for neutrophils. CCL22 was shown to be part of the M2a macrophage repertoire induced after IL-4/IL-13 stimulation [39], with suppression of adaptive immunity in TAMs [6]. CCL22 secretion also enables the attraction of pro-tumor immunosuppressive cells via interactions with CCR4 receptors on the surface of Th2/Treg cells, while autocrine generation of CXCL12 was shown to support tumor growth and metastasis [32], and to modulate the differentiation of monocytes into a phenotype with immunosuppressive and proangiogenic characteristics [93]. Similar to chemokines was the case of the macrophage scavenger receptor M2 marker MSR1 (CD204) that displayed increased spectral counts in the serum/cytokine treated cells, but did not meet the stringent criteria that were utilized for considering protein upregulation. It was interesting, however, to observe the upregulation of C1QBP (Complement C1q binding protein), a multifunctional protein that is involved in mediating and controlling a very large number of cellular processes including transcription regulation, RNA processing, CDKN2A-mediated apoptosis, protein synthesis in mitochondria, mitochondrial OXPHOS, adhesion and chemotaxis. This protein can be found in various cell compartments, and when acting as a receptor, it can bind C1q molecules to inhibit the activation of the complement C1 complex, and therefore immune reactions [32]. Its ability to interact with viral and bacterial proteins, makes it a critical regulator of anti-bacterial and anti-viral inflammatory responses [94].

Altogether, the analysis of the immune signaling cluster revealed that the cells exhibited a strong response to the cytokine treatment, manifesting mainly a tumor-supportive M2a/c phenotype that displayed pronounced collagen deposition and ECM remodeling processes, type II inflammatory responses, production/upregulation of angiogenic factors and processes (FGF1, PDGFRB, VGFR1), upregulation of protease inhibitors (SERPINE1), suppression of adaptive immunity (CCL22), expression of tumor growth and survival chemokines CXCL12 but not of typical inflammatory cytokines (IL-1, IL-6, IL-12, TNF-α) [6], upregulation of scavenger markers (MSR1, LRP1), and display of mostly endocytic but not phagocytic activities. Pro-inflammatory components (MYD88, AIMP1) were observable. Anti-proliferative effects induced by TGFβ and possibly STAT1 were clearly manifested, as evidenced by the analysis of cell cycle regulating components (see below). The impact of adding CCL2 to the stimulating cocktail of cytokines was not clear. On one hand, CCL2 can elicit immunosuppressive effects and support tumor progression by promoting increased invasion through angiogenesis [95], while on the other, it can induce the production of pro-inflammatory cytokines in macrophages to augment their survival [96].

#### Cell cycle regulation

On a background of response to stress, a cluster of functionally altered proteins were representative of various aspects of cell cycle regulation, in particular negative regulation, check points, G1/S or G2/M transition, and apoptotic processes (**Figure 10D**). Representative examples included proteins with functional roles in cell cycle arrest (cycD-CDK4/6 complex inhibitors CDKN1A/CDKN2A), proliferation arrest (CEBPA), G1-to-S phase negative regulation (GADD45GIP1), negative regulation of growth (PML, ILK, TSG101, PHB), transcription activation inhibition (TMF1), epigenetic repression (HDAC2), general RNA polymerase II transcription inhibition (HEXIM1), and regulation of apoptotic processes (PML, SP100, STAT1/STAT3, HTRA2). CDK4 and CDK6 were found upregulated in the cytoplasmic fractions, upholding previous results that showed that cycD-CDK 4 complexes are confined to the cytoplasm, and translocate to the nucleus to regulate cell cycle progression only when the nucleus is large enough [97]. Proteins with roles in regulating Ser/Thr kinase activity, mainly MAPK cascade proteins, directed the cellular responses involved in cell cycle control. Many cell cycle proteins were also common to the previously described immune response and cytokine signaling processes (STAT3, MYD88, TAB1), cytoskeletal remodeling, adhesion, motility, localization, nucleocytoplasmic transport, and viral response. Shared components of a complex network of signaling pathways provided the scaffold for communication between these biological processes (i.e., MAPK, RAF/MAP, PI3K/AKT, c-MET, VEGF, PTEN, Notch, NFKB, Wnt, CLR, TCR and B cell receptor pathways). Proteasomal degradation, as represented by various 26S proteasome subunits, proteasome and adhesion regulating protein (ADRM1), ubiquitin-protein ligase (WWP1) and ubiquitin conjugating enzymes, was common to most signaling pathways. The upregulated proteasome components were associated with the nuclear fraction, mirroring prior findings that have shown that proteasome recruitment at the nuclear pore complex facilitates the regulation of transcription [98] and cell cycle progression by catalyzing the degradation of key cell cycle regulatory proteins. The above processes were expected to occur as a result of TGFβ activities that induce G1/S cell cycle arrest via the expression of cycD/E-CDK4/6 inhibitors, and by making use of the ubiquitin-proteasome pathway [99]. STAT signaling activated by cytokines has been also described to upregulate the expression of proteasomal subunits [100], and IL-4 induced STAT1 activation was shown to result in cell growth inhibition [101].

### Downregulated processes in serum-depleted/cytokine-treated cells

The downregulated processes in the serum-depleted cells related to gene expression, i.e., transcription initiation by RNA polymerase II Mediator, cell cycle G1/S and G2/M transition and checkpoints, chromatin organization, chromosome segregation, cell division, DNA repair, and associated metabolic processes (**Figures 8A** and **9B**). This was evident from the genes and proteins that control G1/S or G2/M transition [e.g., G1/S and G2/M transition cyclin A2 (CCNA2), G2/M-specific cyclin B1 (CCNB1), G1/S and G2/M control CDK (CDK1), key regulator kinases of mitosis (PLK1, and AURKB/AURKC)], spindle assembly, DNA repair and replication, and cytoskeleton organization. The simultaneous upregulation of HEXIM1 (a general RNA polymerase II transcription inhibitor) and of the p16 (CDKN2A) and p21 (CDKN1A) inhibitors of cycD-CDK4/6 and cycE-CDK2 complexes, that mediate G1 cell cycle arrest in response to stress (serum deprivation in this case), corroborated these results. The cytokine treated/serum-depleted cells also had the highest proportion of G1 cells, which was anticipated based on earlier reports that have shown that TGFβ and IL-10 induce anti-proliferative effects and an anti-inflammatory response in microglia cells [102,103]. TGFβ has been also shown to suppress cell cycle progression [50], and mediate cell cycle arrest in G1 by the induction of CDK inhibitors [99]. The lower abundance of cyclin A2 (CCN2A), of G2-specific cyclin B1 (CCNB1), and of the nuclear proliferation marker MKI67 in the cytokine-treated cells was supportive of this result. There were no enriched biological processes specific to immune responses in the protein set with downregulated activity or expression. Rather, some proteins with various metabolic and cell-cell communication functions (kinases, phosphatases) implicated in processes such as transcription, cell cycle progression, ubiquitination, transport and cell junction, had associations with immune response processes. For example, several downregulated cell adhesion (VCAM1), cytoskeleton organization, transport and motility proteins included components with known chemotactic or phagocytic activity (e.g., SRC, ELMO2, PTPRG). In addition, the downregulation of a transcription factor regulating the expression of some immune and inflammatory response genes (CEBPB), and of a scaffold protein involved in transcription regulation and innate immunity (PQBP1), was also observed. Similar downregulated processes were observable in the serum-released cells, as well, advocating overall for the multiple roles that TGFβ plays in inhibiting cell growth and immune responses.

### Upregulated biological processes in serum/cytokine-treated cells

Quantitative profiling of the nuclear and cytoplasmic fractions of the serum/cytokine-treated cells resulted in 715 proteins representing biological processes similar to the ones described for the serum-depleted cells. Correlated, however, with the addition of FBS, the aberrant increase in ribosome biogenesis, one of the most energetically expensive biological processes [104], was unique to the serum-supplemented cells. **Figures 8B** and **9C** depict these changes, and relevant findings are discussed in the followings.

Mitochondrial expression, oxidative phosphorylation and ECM remodeling processes were represented mostly by the same proteins as in the serum-depleted cells. Two mitochondrial protein hubs emerged, one comprising mitochondrial ribosome proteins matched to mitochondrial gene expression and translation, and the other to mitochondrial organization, oxidative phosphorylation, ATP synthesis coupled proton transport, and respiratory electron transport. As most mitochondrial proteins are encoded by nuclear DNA and then translated on cytosolic ribosomes, the mitochondrial inner membrane (TIMM) and outer membrane (TOMM) translocase complexes [105] that import proteins into mitochondria were correspondingly upregulated. An increased expression of collagens, integrins and ECM molecules that participate in cell adhesion and migration correlated with increased integrin signaling [106]. In addition, three noteworthy proteins, peroxidasin (PXDN) which is involved in ECM synthesis, alpha-2-macroglobulin (A2M) which is an inhibitor of proteases such as collagenase and of inflammatory cytokines, and disintegrin/metalloproteinase domain-containing protein 17 (ADAM17) which is responsible for the proteolytic release of cell surface proteins, were observed. Altogether, the ECM cluster was intertwined with membrane organization, cytoskeletal rearrangement, receptor proteins, and members of various signaling cascades. The induction of TGFBI and TGFB1I1 adhesion and migration modulators was observable, and downstream effects of TGFβ signaling with roles in regulating actin cytoskeleton dynamics, cell migration and angiogenesis were manifested via the upregulated RHOA, ROCK1, and SMAD2 (as well as its coactivator SNW1) proteins. By entrapping growth factors, nutrients and cytokines, the extracellular milieu clearly sustained a supportive environment for relaying signals to and inside the cell.

### Immune response

The immune response clusters were less distinct in the serum/cytokine treated cells, most likely due to the dominating cell-cycle related processes induced by the addition of serum. Nevertheless, proteins involved in signal transduction pathways that link the cell membrane receptors to adhesion processes, cytoskeleton organization, transport/localization and intracellular signaling (MAPK, WNT, NFKB, TGFβ, STAT, cell cycle), could be associated with antiviral innate and interferon type I responses (NFKB1, TRAF3, PML, SP100, STAT1/2, ISG15/20, MX1/2, IFIT1, SAMHD1, HLA-A, HLA-C, TRIMs). Intermingled with cell cycle and nuclear import/export processes, notable was the higher level of interferon signaling that was evident from the activation of the interferon-induced proteins (IFIT1, ISG15, ILF3, MX1, MX2, TRIM22). The signal transducers STAT1 and STAT2 which mediate IFN-γ signaling were also present with increased spectral counts in the cells. At the same time, however, PARP14, an ADP-ribosyltransferase which has been proposed to have roles in suppressing pro-inflammatory IFN- γ/STAT1 macrophage activation, while also promoting anti-inflammatory activities via IL-4/STAT6 modulation [107], was upregulated, as well. The protein was also shown to upregulate MRC1 in response to IL-4 stimulation, but MRC1 expression was not detectable in the cells. HLA-C class I histocompatibility antigens, previously detected in the serum depleted cells, were complemented here by HLA-A antigens (A-24α and A-31α) and by a downregulated HLA-DOA, which is a modulator of HLA class II histocompatibility antigen presentation pathways [32]. Its expression is inhibited by IL-10 [11].

The multifaceted NFKB signaling enterprise was also better represented in the serum/cytokine-treated cells, as evidenced by the upregulation of TRAF3 (TNF receptor associated factor)-a protein that is involved in NFKB activation, of NFKB1 (p105 or p50)-which promotes the transcription of inflammatory genes, and of its inhibitor (NFKBIE)-which prevents it from translocating into the nucleus and activating transcription. NFKB1 and NFKBIE were both present in higher abundance in the cytoplasmic fraction of stimulated cells, and NFKBIE in the nucleus too. The common heterodimeric complex partner of NFKB1, RELA or p65, was slightly upregulated in the cytoplasmic fractions, without passing though the 2-fold change criteria. NFKB1 is activated in response to a variety of stimuli and can act as both a transcriptional activator or repressor as a function of the type of complexes that it forms. Previous reports suggest that the ECM molecules laminin or fibronectin, along with certain phagocytic processes, can also trigger NFKB activation via integrin-mediated signaling [108]. However, the NFKB1/RELA complex could not be detected in the nucleus, suggesting that the pathway was not fully activated. Nonetheless, REL (or c-REL), which is one of the transcription factors for HLA-A [11], could be observed with increased counts in the nuclear fraction of cells. Interesting to note was also that an NFKB-activating protein (NKAP) that induces activation upon inflammatory cytokine stimulation, was depleted in the nuclear fractions of both G1 and S cytokine-treated cells (passing the statistical filtering criteria only in G1).

Despite the observed alterations in these pathways, the production of classical pro-inflammatory cytokines did not occur. Only one TNF protein, C1QTNF1 (Complement C1q tumor necrosis factor-related protein 1), was detected with increased counts. The upregulation of several other proteins with roles in immune defense against pathogens (C1QBP, NLRC4, ANKRD17) is worthy, however, of additional comments. Relevant to the findings, the multifunctional C1q complement binding protein C1QBP which was also upregulated in the serum-depleted cells, is a protein that has roles in driving ribosome biogenesis, protein synthesis in mitochondria, and assembly of the mitoribosome. NLRC4 (NLR family CARD domain-containing protein 4), a component of the inflammasome, induces caspase-1 activation and cytokine production in response to pathogens, and ANKRD17 (Ankyrin repeat domain 17) mediates innate antibacterial pathways via NOD1/NOD2 pattern recognition responses [32].

Altogether, while elements of interferon and NFKB signaling were observed in the serum-depleted cells, the molecular differences brought about by the addition of serum appeared to have contributed to augmenting these processes in the serum/cytokine-treated cells. Either canonical NFKB signaling activated by TLR stimulation leading to the expression of certain uncharacteristic IFN signaling and mediator proteins (TLR 3 and 7 were upregulated in the cells), or autocrine signaling through cytokine receptors followed by activation, may have resulted ultimately in a spectrum of microglia responses. Inhibitory effects induced by the anti-inflammatory cytokines, lack of a strong enough signaling input to meet the necessary thresholds, negative feedback loops or cross-talk with other signaling pathways, appeared to have antagonized ultimately a full activation process.

### Cell cycle and ribosome biogenesis

The upregulated cell cycle processes were sustained by proteins indicative of G1/S transition and DNA replication initiation (DNA replication licensing factors MCM-minichromosome maintenance proteins 2-7), DNA primase PRIM1, mitotic prometaphase proteins (chromosome alignment/segregation centromere-interacting proteins, kinetochore function, mitotic checkpoint ZWILCH and KNTC1), components of the nucleopore complex (nucleoporins, AHCTF1), and RNA polymerase transcription initiation/termination proteins (RNA polymerase II subunits, GTFs 2F/2E/2H, ERCC3). Cyclin H (CCNH)-an activator of CDK1/2/4/6, and cyclin Y (CCNY)-a positive regulator of Wnt signaling pathway, were both present with increased counts in the nuclear fractions of cytokine treated cells. Many of the above proteins have roles in both positive and negative regulation of gene expression, and the presence of negative regulators of cell cycle (PML, WEE2, E2F7, CTNNB1, PPP2R5B, ILK, THBS1, EZH2), TP53 activity regulators (PML, TP53RK), and apoptotic proteins (AIFM1, MST4, STK24, MTCH1, SH3GLB1), underscored complex functional, and possibly opposite trends in the dataset. Some cell cycle proteins were also representative of immune/cytokine response and antiviral processes. For example, SAMHD1 which is a regulator of DNA end resection at stalled replication forks is also involved in antiviral defense responses, and the interferon-induced antivirals GTPases MX1 and MX2 have roles in cell death and cell-cycle progression, respectively.

Ribosome biogenesis is considered a hallmark of cell growth and division. The nuclear machinery that mediates DNA transcription and ribosome synthesis prior to mRNA translation was heavily represented by general transcription factors (GTFs), RNA polymerase II (POLR2s), nucleolar ribosome biogenesis proteins (UTPs), nucleolar regulator of RNA polymerase I (NOLC1), and nucleolar proteins involved in pre-rRNA (NOL11, NOP14, NSA2, WD repeat proteins) and rRNA processing (RRPs). The cytoplasmic translation initiation factors (EIF1AX, EIF2B2, EIF2D) complemented this group. It is known that anabolic processes such as ribosome biogenesis and protein synthesis are driven by mTORC signaling in growing cells [103]. It has also been shown that mTORC1 negatively regulates autophagy, and positively affects oxidative metabolism through changes in mitochondrial DNA content [109]. Along these lines of evidence, the activation of two mediators of the mTOR pathway, RPTOR and LAMTOR1, which upon sensing nutrients activate mTORC1 and promote cell growth, was observed. In response to IL-4 stimulation, LAMTOR1, a component of the Ragulator complex, was also shown to be essential for suppressing inflammatory innate immune responses and polarizing macrophages towards an M2 phenotype [110]. As shown, however, by the cell-cycle protein clusters that were downregulated in the cells, and that were indicative overall of impaired cell proliferation (see below), the overexpression of ribosome biogenesis proteins appeared to be rather related to mitochondrial expression, ECM and migratory related processes than to cell cycle progression.

### Downregulated biological processes in serum/cytokine-treated cells

Similar to the serum-depleted cells, the serum/cytokine-treated cells showed a downregulation of processes involved gene expression, transcription regulation, cell cycle, DNA repair, vesicle transport/intracellular trafficking, and cytoskeleton organization (**Figures 8B** and **9D**). Representative proteins included mainly transcription initiation factor TFIID subunits, general transcription factors (GTFs), Mediator subunits of RNA polymerase II transcription, RNA polymerase subunits (mainly I), regulators and activators of transcription (NFIB-Nuclear factor IB, HMGB2, RPS6KA5, BCLAF1, and THRAP 3), replication factor RFC1, and components of the E3 ubiquitin-protein ligase complexes (APC, BCR, SCF) that mediate the ubiquitination and degradation of target proteins (cyclins, CDKs), thereby controlling cell cycle G1/S transition and exit from mitosis (ANAPC1/10, CUL3, SKP2, KLHL9). Relevant to note, however, that BCLAF1, THRAP3 and SNIP1 are part of a complex SNARP that in addition to roles in transcription, also regulates CCND1 (cyclin D1) mRNA stability. Unlike BCLAF1 and THRAP3 that were downregulated in the serum-cytokine treated cells, SNIP1 was not. The levels of SNIP1 increase after serum stimulation and promote cell cycle progression from G1 to S by positively regulating cyclin D1 expression. Cyclin D1 was not detectable, but the downregulation of cyclin A2 (CCNA2) and B2 (CCNB1), as well as of cyclin T2 (CCNT2)-a subunit of the positive elongation transcription factor, confirmed the suppressed status of these cells. Further, the identification with lower nuclear counts of CDT1, a DNA replication licensing factor with roles in generating the pre-replication complex, was indicative of its possible degradation and impaired proliferation. Moreover, PLK1 (Polo-like kinase), a Ser/Thr kinase which increases in expression level during mitosis, was downregulated in the nuclear fractions of cytokine-treated cells, also suggesting repressed proliferation. In the cytoplasmic fraction, the counts were slightly increased, but overall low.

MHCII class antigen presentation was affected by the downregulation of HLA-DOA, vesicle-mediated protein transport, and microtubule-dependent and Rho-mediated signaling. Downregulated proteins involved in adhesion, cytoskeletal re-arrangement, motility/migration (ENG, PIK3CB), phagocytosis (ELMO2, DOCK1), as well as in intracellular membrane/vesicle trafficking and endocytosis (RAB13, RAB5A, DNM1, DNM3, kinesins), impacted processes related to chemotaxis and phagocytosis. On a global level, changes in the activity of proteins with roles in the transcription regulation and activation of inflammatory genes (e.g., RPS6KA5, a Ser/Thr kinase required for regulating the expression of transcription factors STAT3 and NFKB subunit RELA), in the activation of signaling cascades involved in cell proliferation/survival and motility (PIK3CB-a catalytic subunit of PI3K), in proliferation/migration and PRR signaling (HMGB2), TGFβ signaling (ENG and SMAD4, with ENG being a TGFβ co-receptor that leads to the activation of SMAD transcription factors, also having roles in regulating angiogenesis), and in cell-cell recognition and adhesion processes (VCAM1), are expected to affect innate immunity and inflammatory responses. Overall, the results confirm the role of TGFβ in cell cycle arrest, cytokinesis, suppression of immune responses, and the development of a cancer-supportive microglia phenotype. Changes in the activity of CD33 (Myeloid cell surface antigen) that inhibits immune cell activation [57], and that requires PI3K to exert its suppressive effects, suggest the presence of both immune-activating and deactivating trends. It was interesting to note the downregulation of TLK2, a Ser/Thr kinase with roles in chromatin assembly and transcription, that is also a negative regulator of autophagy induced by amino acid starvation [32].

### Tumorigenic phenotype of microglia activated by the anti-inflammatory cytokines

The protein networks and signaling pathways that were overexpressed in the cytokine-treated HMC3 cells uncovered numerous mechanisms by which the microglia cells can support tumor progression in the brain. These included dynamic changes in the composition of the ECM that foster tumor attachment and invasion, increased expression of proteins that facilitate angiogenesis, secretion of cytokines and growth factors that promote tumor growth, and expression of proteins that support the recruitment of cells that suppress immune activation. In **Figure 11**, we propose a model for the molecular mechanisms that operate in microglia cells to regulate their pro-tumorigenic functions. The key determinants of the tumor-supportive microglia phenotype, stimulated by IL-4, IL-13, IL-10, TGFβ and CCL2, were the proteins of the MAPK, STAT, TGFβ, NFKB, PI3K, mTOR and integrin signaling pathways. A pivotal outcome of these activated networks was the remodeling of the ECM matrix, as evidenced by increased collagen deposition, stiffening of collagen fibers by enzymes such as LOX and SPARC, and production of proteases (CTSBB, CTSD) and protease inhibitors (SERPINE1) that alter the intercellular matrix composition to ultimately support tumor adhesion and invasion. Together with collagen, additional matrix proteins such as FN1, VTN, FBN trap growth factors and cytokines to facilitate the transmission of signals to- and within the cells to trigger the expression of genes that control protein synthesis, cell survival, migration, and angiogenic processes implicated in cancer progression. Secreted proteases enable the release of growth factors and signaling peptides from the ECM, while SERPINE1 not only regulates cell adhesion and spreading, but also functions by enhancing angiogenesis and by recruiting M2 polarized macrophages or microglia to the site of tumorigenesis [111]. TNC favors tumor growth and invasion by also affecting the matrix’s tensile strength, as well as by suppressing immune activation by limiting the infiltration of lymphocytes in the brain tumors [112]. Collagen-induced DDR receptors in cancer initiate pro-migratory and invasion programs [72, 113], and together with other growth factor receptors, cell adhesion and ECM remodeling molecules facilitate metastatic processes. Integrin-mediated signaling has diverse roles in supporting tumor progression via processes that range from angiogenesis to adhesion, survival, motility and microglial recruitment [73,74,106]. By bridging the ECM with the intracellular environment, integrin receptors cause changes to the actin cytoskeleton [76] to promote the migration of cells in the surrounding environment.

**Figure 11.**
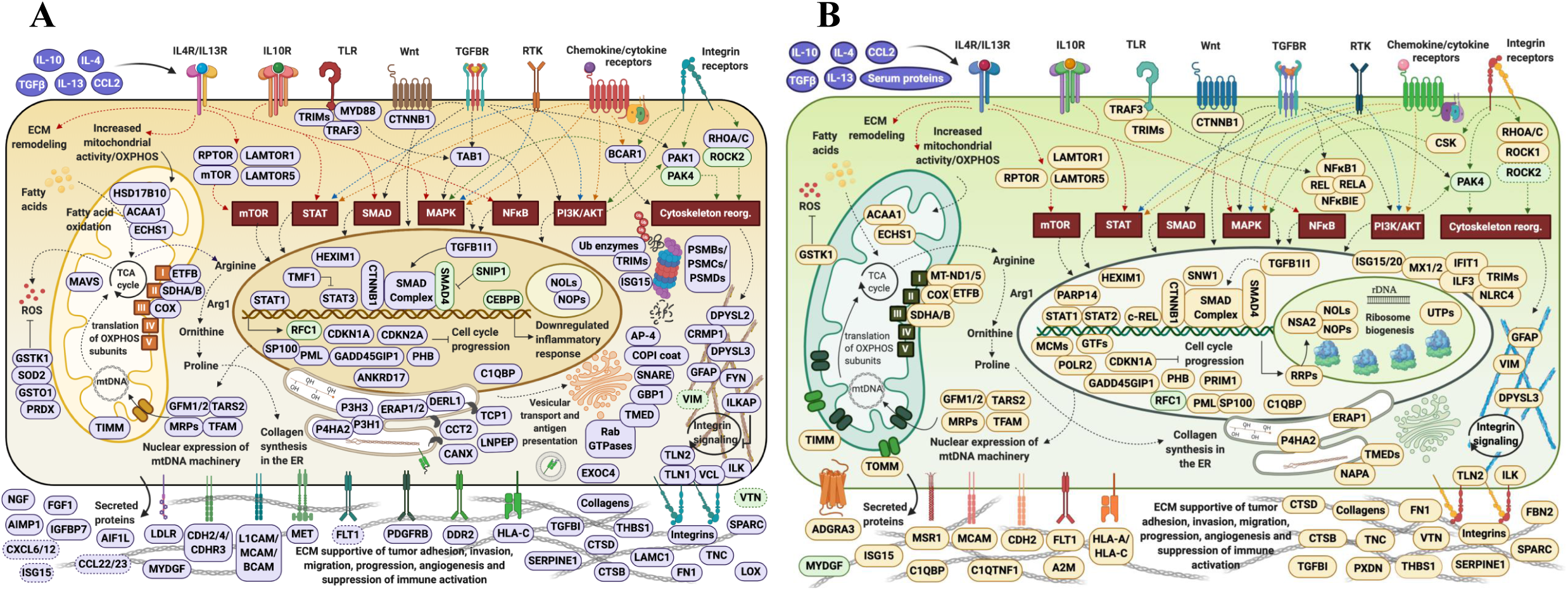
Proposed model for upregulated molecular mechanisms that operate in pro-tumorigenic HMC3 cells. (**A**) Cytokine-treated/serum-depleted cells. (**B**) Cytokine/serum-treated cell. Notes: Genes were placed in the cellular compartment in which they exert their main function. Genes highlighted in purple or amber displayed increased spectral counts, while genes highlighted in green displayed decreased spectral counts in the cytokine treated cells.

The immunosuppressive and tumorigenic characteristics of microglia were further augmented by the secretion of growth factors such as NGF which counters the inflammatory responses mounted by microglia cells [114], MYDGF which increases viability of cancer cells [115], FGF1 which supports angiogenesis, tumor growth and invasion, and of cytokines CCL22 and CXCL12 which attract pro-tumor immunosuppressive cells and support metastasis [6]. Through an increased expression of MHC class I molecule, tumor-associated macrophages and microglia have demonstrated diminished immune activation against tumor-cells and induced expression of angiogenic factors and cytokines [116]. Moreover, the anti-tumor inflammatory activation of microglia was directly affected by reduced cell proliferation that was linked to TGFβ and IL-10 signaling mechanisms. Altogether, the upregulated biological processes in HMC3 suggest that the microglia cells favor the formation of a tumor-supportive niche when they are subjected to an anti-inflammatory microenvironment *in-vitro*.

## Conclusions

Proteomic analysis of HMC3 cells treated with anti-inflammatory cytokines exposed a vast array of altered biological and immunological functions. These processes distinguished themselves more clearly in the serum-depleted cells. Altogether, the combined effect of IL-4, IL-13, IL-10, TGFβ and CCL2 resulted in the induction of a spectrum of M2a/c microglia phenotypes with capabilities for inducing matrix deposition, ECM-remodeling and angiogenic processes, and overall, a tumor-supportive niche that facilitates growth and invasion. Mitochondrial gene expression and respiration were highly upregulated processes, resulting in altered arginine metabolism and collagen synthesis. As evidenced by the upregulation of certain receptors, cytokines and endocytic activities, the stimulated cells were inclined to rather exhibit an immunosuppressive response, even though pro-inflammatory defense elements were observable. MAPK signaling was central to all cell communication pathways, and STAT components were involved in mediating several biological outcomes. The negative impact of TGFβ on cell cycle progression was clearly manifested through the altered activity of proteins involved in G1/S and G2/M transition, gene expression, cell division, and apoptosis. NFKB signaling manifested itself mainly in the serum/cytokine-treated cells but it did not emerge as being fully activated. It was accompanied, however, by several proteins representative of interferon signaling. Altogether, the study underscores the complex network of interactions that are activated in stimulated microglia cells, and demonstrates the benefits of proteomic profiling for gaining insights into the many, still unknown, environment-contingent mechanisms that drive cancer progression. The landscape that underpins the range of immunosuppressive activities triggered by anti-inflammatory cytokines will point to possible novel therapeutic targets enabled by co-operative microglia cells, and will be particularly useful to guiding future drug discovery efforts.

## Data availability

The mass spectrometry raw files were deposited to the ProteomeXchange Consortium via the PRIDE partner repository with the following dataset identifiers: PXD023163 and PXD023166.

## Acknowledgment

This work was supported by an award from the National Institute of General Medical Sciences (Grant No. 1R01GM121920) to IML.

## Author contributions

SA performed the experiments; SA and IML analyzed the data and wrote the manuscript; IML coordinated the work. All authors reviewed and approved the final version of the manuscript.

## References

1. Nakajima, K., and Kohsaka, S. (1993). Functional roles of microglia in the brain. Neurosci. Res. 17(3), 187–203.

2. Nimmerjahn, A., Kirchhoff, F., and Helmchen, F. (2005). Resting microglial cells are highly dynamic surveillants of brain parenchyma in vivo. Science 308(5726), 1314–1318.

3. Hanisch, U. K., and Kettenmann, H. (2007). Microglia: active sensor and versatile effector cells in the normal and pathologic brain. Nat. Neurosci. 10(11), 1387–1394.

4. Roesch, S., Rapp, C., Dettling, S., and Herold-Mende, C. (2018). When immune cells turn bad—tumor-associated microglia/macrophages in glioma. Int. J. Mol. Sci. 19(2), 436.

5. Wu, S. Y., and Watabe, K. (2017). The roles of microglia/macrophages in tumor progression of brain cancer and metastatic disease. Front. Biosci. (Landmark edition) 22, 1805.

6. Sica, A., Schioppa, T., Mantovani, A., and Allavena, P. (2006). Tumour-associated macrophages are a distinct M2 polarised population promoting tumour progression: potential targets of anti-cancer therapy. Eur. J. Cancer 42(6), 717–727.

7. Zhang, J., Sarkar, S., Cua, R., Zhou, Y., Hader, W., and Yong, V. W. (2012). A dialog between glioma and microglia that promotes tumor invasiveness through the CCL2/CCR2/interleukin-6 axis. Carcinogenesis 33(2), 312–319.

8. Wang, Q., He, Z., Huang, M., Liu, T., Wang, Y., Xu, H., et al. (2018). Vascular niche IL-6 induces alternative macrophage activation in glioblastoma through HIF-2α. Nat. Commun. 9(1), 1–15.

9. Cherry, J. D., Olschowka, J. A., and O’Banion, M. K. (2014). Neuroinflammation and M2 microglia: the good, the bad, and the inflamed. J. Neuroinflamm. 11(1), 98.

10. Li, W., and Graeber, M. B. (2012). The molecular profile of microglia under the influence of glioma. Neuro-oncology 14(8), 958–978.

11. Cruz-Tapias, P., Castiblanco, J., and Juan-Manuel, A. (2013). “Major histocompatibility complex: antigen processing and presentation,” in Autoimmunity: From Bench to Bedside, eds J. M. Anaya, Y. Shoenfeld, and A. Rojas-Villarraga, (Bogota: El Rosario University Press), 169–184.

12. Forrester, J. V., McMenamin, P. G., and Dando, S. J. (2018). CNS infection and immune privilege. Nat. Rev. Neurosci. 19(11), 655–671.

13. Alexandru, D., Bota, D. A., and Linskey, M. E. (2012). “Epidemiology of central nervous system metastases,” in Current and Future Management of Brain Metastasis, eds Kim D.G. and Lunsford L.D., (Prog Neurol Surg. Basel, Karger), 25, 13–29.

14. Cagney, D. N., Martin, A. M., Catalano, P. J., Redig, A. J., Lin, N. U., Lee, E. Q., et al. (2017). Incidence and prognosis of patients with brain metastases at diagnosis of systemic malignancy: a population-based study. Neuro-oncology 19(11), 1511–1521.

15. You, H., Baluszek, S., and Kaminska, B. (2019). Immune microenvironment of brain metastases-are microglia little helpers? Front. Immunol. 10, 1941.

16. Choi, J., Mai, N., Jackson, C., Belcaid, Z., and Lim, M. (2018). It takes two: potential therapies and insights involving microglia and macrophages in glioblastoma. Population 10, 11.

17. Cheray, M., and Joseph, B. (2018). Epigenetics control microglia plasticity. Front. Cell. Neurosci. 12, 243.

18. Kolosowska, N., Keuters, M. H., Wojciechowski, S., Keksa-Goldsteine, V., Laine, M., Malm, T., et al. (2019). Peripheral administration of IL-13 induces anti-inflammatory microglial/macrophage responses and provides neuroprotection in ischemic stroke. Neurotherapeutics 16(4), 1304–1319.

19. Fitzgerald, D. P., Palmieri, D., Hua, E., Hargrave, E., Herring, J. M., Qian, Y., et al. (2008). Reactive glia are recruited by highly proliferative brain metastases of breast cancer and promote tumor cell colonization. Clin. Exp. Metastasis 25(7), 799–810.

20. Janabi, N., Peudenier, S., Héron, B., Ng, K. H., and Tardieu, M. (1995). Establishment of human microglial cell lines after transfection of primary cultures of embryonic microglial cells with the SV40 large T antigen. Neurosci. Lett. 195(2), 105–108.

21. Russo, C. D., Cappoli, N., Coletta, I., Mezzogori, D., Paciello, F., Pozzoli, G., et al. (2018). The human microglial HMC3 cell line: where do we stand? A systematic literature review. J. Neuroinflamm. 15(1), 259.

22. Vergara, D., Nigro, A., Romano, A., De Domenico, S., Damato, M., Franck, J., et al. (2019). Distinct Protein Expression Networks are Activated in Microglia Cells after Stimulation with IFN-γ and IL-4. Cells 8(6), 580.

23. Chiavari, M., Ciotti, G. M. P., Navarra, P., and Lisi, L. (2019). Pro-inflammatory activation of a new immortalized human microglia cell line. Brain Sci. 9(5), 111.

24. Lively, S., and Schlichter, L. C. (2018). Microglia responses to pro-inflammatory stimuli (LPS, IFNγ+ TNFα) and reprogramming by resolving cytokines (IL-4, IL-10). Front. Cell. Neurosci. 12, 215.

25. Deng, J., Julian, M. H., and Lazar, I. M. (2018). Partial enzymatic reactions: A missed opportunity in proteomics research. Rapid Commun. Mass Spectrom. 32(23), 2065–2073.

26. Tenga, M. J., and Lazar, I. M. (2013). Proteomic snapshot of breast cancer cell cycle: G1/S transition point. Proteomics, 13(1), 48–60.

27. Huang, D. W., Sherman, B. T., and Lempicki, R. A. (2009). Bioinformatics enrichment tools: paths toward the comprehensive functional analysis of large gene lists. Nucleic Acids Res., 37(1), 1–13.

28. Szklarczyk, D., Gable, A. L., Lyon, D., Junge, A., Wyder, S., Huerta-Cepas, J., et al. (2019). STRING v11: protein–protein association networks with increased coverage, supporting functional discovery in genome-wide experimental datasets. Nucleic Acids Res.,47(D1), D607–D613.

29. Kanehisa, M., and Goto, S. (2000). KEGG: kyoto encyclopedia of genes and genomes. Nucleic Acids Res., 28(1), 27–30.

30. Jassal, B., Matthews, L., Viteri, G., Gong, C., Lorente, P., Fabregat, A., et al. (2020). The reactome pathway knowledgebase. Nucleic Acids Res., 48(D1), D498–D503.

31. UniProt Consortium. (2019). UniProt: a worldwide hub of protein knowledge. Nucleic Acids Res., 47(D1), D506–D515.

32. Stelzer, G., Rosen, N., Plaschkes, I., Zimmerman, S., Twik, M., Fishilevich, S., et al. (2016). The GeneCards suite: from gene data mining to disease genome sequence analyses. Curr. Protoc. Bioinformatics, 54(1), 1–30.

33. BioRender, https://biorender.com.

34. Berghoff, A. S., Lassmann, H., Preusser, M., and Höftberger, R. (2013). Characterization of the inflammatory response to solid cancer metastases in the human brain. Clin. Exp. Metastasis 30(1), 69–81.

35. Matias, D., Balça-Silva, J., da Graça, G. C., Wanjiru, C. M., Macharia, L. W., Nascimento, C. P., et al. (2018). Microglia/astrocytes–glioblastoma crosstalk: crucial molecular mechanisms and microenvironmental factors. Front. Cell. Neurosci. 12, 235.

36. Panni, R. Z., Linehan, D. C., and DeNardo, D. G. (2013). Targeting tumor-infiltrating macrophages to combat cancer. Immunotherapy 5(10), 1075–1087.

37. Tang, Y., and Le, W. (2016). Differential roles of M1 and M2 microglia in neurodegenerative diseases. Mol. Neurobiol. 53(2), 1181–1194.

38. Zhou, X., Spittau, B., and Krieglstein, K. (2012). TGFβ signalling plays an important role in IL4-induced alternative activation of microglia. J. Neuroinflamm. 9(1), 210.

39. Mantovani, A., Sica, A., Sozzani, S., Allavena, P., Vecchi, A., and Locati, M. (2004). The chemokine system in diverse forms of macrophage activation and polarization. Trends. Immunol. 25(12), 677–686.

40. Könnecke, H., and Bechmann, I. (2013). The role of microglia and matrix metalloproteinases involvement in neuroinflammation and gliomas. Clin. Dev. Immunol. 2013.

41. Rath, M., Müller, I., Kropf, P., Closs, E. I., and Munder, M. (2014). Metabolism via arginase or nitric oxide synthase: two competing arginine pathways in macrophages. Front. Immunol. 5, 532.

42. Ueno, T., Toi, M., Saji, H., Muta, M., Bando, H., Kuroi, K., et al. (2000). Significance of macrophage chemoattractant protein-1 in macrophage recruitment, angiogenesis, and survival in human breast cancer. Clin. Cancer Res. 6(8), 3282–3289.

43. Hinojosa, A. E., Garcia-Bueno, B., Leza, J. C., and Madrigal, J. L. (2011). CCL2/MCP-1 modulation of microglial activation and proliferation. J. Neuroinflamm. 8(1), 77.

44. Réu, P., Khosravi, A., Bernard, S., Mold, J. E., Salehpour, M., Alkass, K., et al. (2017). The lifespan and turnover of microglia in the human brain. Cell Rep. 20(4), 779–784.

45. Graeber, M. B., Streit, W. J., and Kreutzberg, G. W. (1988). The microglial cytoskeleton: vimentin is localized within activated cells in situ. J. Neurocytol. 17(4), 573–580.

46. Jin, M. M., Wang, F., Qi, D. I., Liu, W. W., Gu, C., Mao, C. J., et al. (2018). A critical role of autophagy in regulating microglia polarization in neurodegeneration. Front. Aging Neurosci. 10, 378.

47. Liliensiek, S. J., Schell, K., Howard, E., Nealey, P., and Murphy, C. J. (2006). Cell sorting but not serum starvation is effective for SV40 human corneal epithelial cell cycle synchronization. Exp. Eye Res. 83(1), 61–68.

48. Cooper, S. (1998). Mammalian cells are not synchronized in G1-phase by starvation or inhibition: considerations of the fundamental concept of G1-phase synchronization. Cell Prolif. 31(1), 9–16.

49. Keyomarsi, K., Sandoval, L., Band, V., and Pardee, A. B. (1991). Synchronization of tumor and normal cells from G1 to multiple cell cycles by lovastatin. Cancer Res. 51(13), 3602–3609.

50. Zhang, Y. E. (2017). Non-Smad signaling pathways of the TGF-β family. Cold Spring Harb. Perspect. Biol. 9(2), a022129.

51. Starr, D. A., and Fridolfsson, H. N. (2010). Interactions between nuclei and the cytoskeleton are mediated by SUN-KASH nuclear-envelope bridges. Annu. Rev. Cell Dev. Biol. 26, 421–444.

52. Lee, W., and Lazar, I. M. (2014). Endogenous protein “Barcode” for data validation and normalization in quantitative MS analysis. Anal. Chem. 86(13), 6379–6386.

53. Böttcher, C., Schlickeiser, S., Sneeboer, M. A., Kunkel, D., Knop, A., Paza, E., et al. (2019). Human microglia regional heterogeneity and phenotypes determined by multiplexed single-cell mass cytometry. Nat. Neurosci. 22(1), 78–90.

54. Wei, J., Gabrusiewicz, K., and Heimberger, A. (2013). The controversial role of microglia in malignant gliomas. Clin. Dev. Immunol. 2013.

55. Butovsky, O., Jedrychowski, M. P., Moore, C. S., Cialic, R., Lanser, A. J., Gabriely, G., et al. (2014). Identification of a unique TGF-β–dependent molecular and functional signature in microglia. Nat. Neurosci. 17(1), 131–143.

56. Melief, J., Sneeboer, M. A. M., Litjens, M., Ormel, P. R., Palmen, S. J. M. C., Huitinga, I., et al. (2016). Characterizing primary human microglia: A comparative study with myeloid subsets and culture models. Glia 64(11), 1857–1868.

57. Crocker, P. R., and Varki, A. (2001). Siglecs, sialic acids and innate immunity. Trends Immunol. 22(6), 337–342.

58. Zhang, S., Wu, M., Peng, C., Zhao, G., and Gu, R. (2017). GFAP expression in injured astrocytes in rats. Exp. Ther. Med. 14(3), 1905–1908.

59. Izquierdo, P., Attwell, D., and Madry, C. (2019). Ion channels and receptors as determinants of microglial function. Trends Neurosci. 42(4), 278–292.

60. ElAli, A., and Rivest, S. (2016). Microglia ontology and signaling. Front. Cell Dev. Biol. 4, 72.

61. Liu, H., Leak, R. K., and Hu, X. (2016). Neurotransmitter receptors on microglia. Stroke Vasc. Neurol. 1(2), 52–58.

62. Domercq, M., Vazquez, N., and Matute, C. (2013). Neurotransmitter signaling in the pathophysiology of microglia. Front. Cell. Neurosci. 7, 49.

63. Ghosh, M., Xu, Y., and Pearse, D. D. (2016). Cyclic AMP is a key regulator of M1 to M2a phenotypic conversion of microglia in the presence of Th2 cytokines. J. Neuroinflamm. 13(1), 9.

64. Janda, E., Boi, L., and Carta, A. R. (2018). Microglial phagocytosis and its regulation: a therapeutic target in Parkinson’s disease? Front. Mol. Neurosci. 11, 144.

65. Canton, J., Neculai, D., and Grinstein, S. (2013). Scavenger receptors in homeostasis and immunity. Nat. Rev. Immunol. 13(9), 621–634.

66. Todt, J. C., Hu, B., & Curtis, J. L. (2008). The scavenger receptor SR-AI/II (CD204) signals via the receptor tyrosine kinase Mertk during apoptotic cell uptake by murine macrophages. J. Leukoc. Biol. 84(2), 510–518.

67. Fu, R., Shen, Q., Xu, P., Luo, J. J., and Tang, Y. (2014). Phagocytosis of microglia in the central nervous system diseases. Mol. Neurobiol. 49(3), 1422–1434.

68. Gumienny, T. L., Brugnera, E., Tosello-Trampont, A. C., Kinchen, J. M., Haney, L. B., Nishiwaki, K., et al. (2001). CED-12/ELMO, a novel member of the CrkII/Dock180/Rac pathway, is required for phagocytosis and cell migration. Cell 107(1), 27–41.

69. Yang, I., Han, S. J., Kaur, G., Crane, C., and Parsa, A. T. (2010). The role of microglia in central nervous system immunity and glioma immunology. J. Clin. Neurosci. 17(1), 6–10.

70. Galván-Peña, S., and O’Neill, L. A. (2014). Metabolic reprograming in macrophage polarization. F. Immunol. 5, 420.

71. Orihuela, R., McPherson, C. A., and Harry, G. J. (2016). Microglial M1/M2 polarization and metabolic states. Br. J. Pharmacol. 173(4), 649–665.

72. Xu, S., Xu, H., Wang, W., Li, S., Li, H., Li, T., et al. (2019). The role of collagen in cancer: from bench to bedside. J. Transl. Med. 17(1), 309

73. Fang, M., Yuan, J., Peng, C., and Li, Y. (2014). Collagen as a double-edged sword in tumor progression. Tumor Biol. 35(4), 2871–2882.

74. Winkler, J., Abisoye-Ogunniyan, A., Metcalf, K. J., and Werb, Z. (2020). Concepts of extracellular matrix remodelling in tumour progression and metastasis. Nat. Comm. 11(1), 1–19.

75. Qian, B. Z., and Pollard, J. W. (2010). Macrophage diversity enhances tumor progression and metastasis. Cell 141(1), 39–51.

76. Lampi, M. C., and Reinhart-King, C. A. (2018). Targeting extracellular matrix stiffness to attenuate disease: From molecular mechanisms to clinical trials. Sci. Transl. Med. 10(422), eaao0475.

77. Vidak, E., Javoršek, U., Vizovišek, M., and Turk, B. (2019). Cysteine cathepsins and their extracellular roles: shaping the microenvironment. Cells 8(3), 264.

78. Guo, X., and Wang, X. F. (2009). Signaling cross-talk between TGF-β/BMP and other pathways. Cell Res. 19(1), 71–88.

79. Luo, K. (2017). Signaling cross talk between TGF-β/Smad and other signaling pathways. Cold Spring Harb. Perspect. Biol. 9(1), a022137.

80. Malsin, E. S., Kim, S., Lam, A. P., and Gottardi, C. J. (2019). Macrophages as a source and recipient of Wnt signals. F. Immunol. 10, 1813.

81. Van den Bossche, J., Laoui, D., Naessens, T., Smits, H. H., Hokke, C. H., Stijlemans, B., et al. (2015). E-cadherin expression in macrophages dampens their inflammatory responsiveness in vitro, but does not modulate M2-regulated pathologies in vivo. Sci. Rep. 5, 12599.

82. De Pascalis, C., and Etienne-Manneville, S. (2017). Single and collective cell migration: the mechanics of adhesions. Mol. Biol. Cell. 28(14), 1833–1846.

83. Turaga, S. M., and Lathia, J. D. (2016). Adhering towards tumorigenicity: altered adhesion mechanisms in glioblastoma cancer stem cells. CNS Oncol. 5(4), 251–259.

84. Verma, R., Balakrishnan, L., Sharma, K., Khan, A. A., Advani, J., Gowda, H., et al. (2016). A network map of Interleukin-10 signaling pathway. J. Cell. Commun. Signal. 10(1), 61–67.

85. Voßhenrich, C. A., and Di Santo, J. P. (2002). Interleukin signaling. Curr. Biol. 12(22), R760–R763.

86. Avalle, L., Pensa, S., Regis, G., Novelli, F., and Poli, V. (2012). STAT1 and STAT3 in tumorigenesis: A matter of balance. Jak-stat 1(2), 65–72.

87. Herrero, C., Hu, X., Li, W. P., Samuels, S., Sharif, M. N., Kotenko, S. et al. (2003). Reprogramming of IL-10 activity and signaling by IFN-γ. J. Immunol. 171(10), 5034–5041.

88. Yan, Z., Gibson, S. A., Buckley, J. A., Qin, H., and Benveniste, E. N. (2018). Role of the JAK/STAT signaling pathway in regulation of innate immunity in neuroinflammatory diseases. Clin. Immunol. 189, 4–13.

89. Martinez, F. O., and Gordon, S. (2014). The M1 and M2 paradigm of macrophage activation: time for reassessment. F1000 Prime Rep. 6.

90. Hornbeck, P. V., Zhang, B., Murray, B., Kornhauser, J. M., Latham, V., Skrzypek, E. (2015). PhosphoSitePlus, 2014: mutations, PTMs and recalibrations. Nucleic Acids Res. 43, D512–520.

91. Loureiro, J., and Ploegh, H. L. (2006). Antigen presentation and the ubiquitin-proteasome system in host–pathogen interactions. Adv. Immunol. 92, 225–305.

92. Ferrington, D. A., and Gregerson, D. S. (2012). Immunoproteasomes: structure, function, and antigen presentation. Prog. Mol. Biol. Transl. Sci. 109, 75–112

93. Ruytinx, P., Proost, P., Van Damme, J., and Struyf, S. (2018). Chemokine-induced macrophage polarization in inflammatory conditions. Front. Immunol. 9, 1930.

94. Barna, J., Dimén, D., Puska, G., Kovács, D., Csikós, V., Oláh, S., et al. (2019). Complement component 1q subcomponent binding protein in the brain of the rat. Sci. Rep. 9(1), 1–19.

95. Lim, S. Y., Yuzhalin, A. E., Gordon-Weeks, A. N., and Muschel, R. J. (2016). Targeting the CCL2-CCR2 signaling axis in cancer metastasis. Oncotarget. 7(19), 28697.

96. Gschwandtner, M., Derler, R., and Midwood, K. S. (2019). More than just attractive: how CCL2 influences myeloid cell behavior beyond chemotaxis. Front. Immunol. 10, 2759.

97. Wang, Z., Xie, Y., Zhang, L., Zhang, H., An, X., Wang, T., and Meng, A. (2008). Migratory localization of cyclin D2-Cdk4 complex suggests a spatial regulation of the G1-S transition. Cell Struct. Funct. 0809290029-0809290029.

98. Albert, S., Schaffer, M., Beck, F., Mosalaganti, S., Asano, S., Thomas, H. F., et al. (2017). Proteasomes tether to two distinct sites at the nuclear pore complex. Proc. Natl. Acad. Sci. 114(52), 13726–13731.

99. Zhang, F., Mönkkönen, M., Roth, S., and Laiho, M. (2002). TGF-β induced G1 cell cycle arrest requires the activity of the proteasome pathway. Exp. Cell Res. 281(2), 190–196.

100. Motosugi, R., and Murata, S. (2019). Dynamic regulation of proteasome expression. Front. Mol. Biosci. 6, 30.

101. Chang, T. L. Y., Peng, X., and Fu, X. Y. (2000). Interleukin-4 mediates cell growth inhibition through activation of Stat1. J. Biol. Chem. 275(14), 10212–10217.

102. Suzumura, A., Sawada, M., Yamamoto, H., and Marunouchi, T. (1993). Transforming growth factor-beta suppresses activation and proliferation of microglia in vitro. J. Immunol. 151(4), 2150–2158.

103. O’Farrell, A. M., Liu, Y., Moore, K. W., and Mui, A. L. F. (1998). IL-10 inhibits macrophage activation and proliferation by distinct signaling mechanisms: evidence for Stat3-dependent and-independent pathways. The EMBO J. 17(4), 1006–1018.

104. Mayer, C., and Grummt, I. (2006). Ribosome biogenesis and cell growth: mTOR coordinates transcription by all three classes of nuclear RNA polymerases. Oncogene. 25(48), 6384–6391.

105. Chacinska, A., Koehler, C. M., Milenkovic, D., Lithgow, T., and Pfanner, N. (2009). Importing mitochondrial proteins: machineries and mechanisms. Cell. 138(4), 628–644.

106. Lu, P., Weaver, V. M., and Werb, Z. (2012). The extracellular matrix: a dynamic niche in cancer progression. J. Cell Biol. 196(4), 395–406.

107. Iwata, H., Goettsch, C., Sharma, A., Ricchiuto, P., Goh, W. W. B., Halu, A, et al. (2016). PARP9 and PARP14 cross-regulate macrophage activation via STAT1 ADP-ribosylation. Nat. Comm. 7(1), 1–19.

108. Tong, L., and Tergaonkar, V. (2014). Rho protein GTPases and their interactions with NFκB: crossroads of inflammation and matrix biology. Biosci. Rep. 34(3).

109. Laplante, M., and Sabatini, D. M. (2012). mTOR signaling in growth control and disease. Cell. 149(2), 274–293.

110. Kimura, T., Nada, S., Takegahara, N., Okuno, T., Nojima, S., Kang, S., et al. (2016). Polarization of M2 macrophages requires Lamtor1 that integrates cytokine and amino-acid signals. Nat. Comm. 7(1), 1–17.

111. Kubala, M. H., Punj, V., Placencio-Hickok, V. R., Fang, H., Fernandez, G. E., Sposto, R., and DeClerck, Y. A. (2018). Plasminogen activator inhibitor-1 promotes the recruitment and polarization of macrophages in cancer. Cell Rep. 25(8), 2177–2191.

112. Huang, J. Y., Cheng, Y. J., Lin, Y. P., Lin, H. C., Su, C. C., Juliano, R., and Yang, B. C. (2010). Extracellular matrix of glioblastoma inhibits polarization and transmigration of T cells: the role of tenascin-C in immune suppression. J. Immunol. 185(3), 1450–1459.

113. Valiathan, R. R., Marco, M., Leitinger, B., Kleer, C. G., and Fridman, R. (2012). Discoidin domain receptor tyrosine kinases: new players in cancer progression. Cancer Metastasis Rev. 31(1-2), 295–321.

114. Fodelianaki, G., Lansing, F., Bhattarai, P., Troullinaki, M., Zeballos, M. A., Charalampopoulos, I., et al. (2019). Nerve Growth Factor modulates LPS-induced microglial glycolysis and inflammatory responses. Exp Cell Res. 377(1-2), 10–16.

115. Oliveira, A. I., Anjo, S. I., de Castro, J. V., Serra, S. C., Salgado, A. J., Manadas, B., and Costa, B. M. (2017). Crosstalk between glial and glioblastoma cells triggers the “go-or-grow” phenotype of tumor cells. Cell Commun. Signal. 15(1), 1–12.

116. Marchesi, M., Andersson, E., Villabona, L., Seliger, B., Lundqvist, A., Kiessling, R., and Masucci, G. V. (2013). HLA-dependent tumour development: a role for tumour associate macrophages? J. Transl. Med. 11(1), 247.

