## Supplemental_table_2 for "Systems-Level Proteomics Evaluation of Microglia Response to Tumor-Supportive Anti-inflammatory Cytokines"

**Supplemental Table 2: UniProt ID, gene abbreviations, and description**

| **Uniprot ID** | **Gene name** | **Gene description** |
| --- | --- | --- |
| P01023 | A2M | Alpha-2-macroglobulin |
| P49588 | AARS2 | Alanine--tRNA ligase, mitochondrial |
| P09110 | ACAA1 | 3-ketoacyl-CoA thiolase, peroxisomal |
| P78536 | ADAM17 | Disintegrin and Metalloproteinase Domain-Containing Protein 17 |
| Q14246 | ADGRE1 | Adhesion G protein-coupled receptor E1 |
| Q16186 | ADRM1 | Proteasomal ubiquitin receptor ADRM1 |
| Q8WYP5 | AHCTF1 | Protein ELYS |
| P55008 | AIF1 | Allograft inflammatory factor 1 |
| Q9BQI0 | AIF1L | Allograft inflammatory factor 1-like |
| O95831 | AIFM1 | Apoptosis-inducing factor 1, mitochondrial |
| Q12904 | AIMP1 | Aminoacyl tRNA synthase complex-interacting multifunctional protein 1 |
| P49419 | ALDH7A1 | Alpha-aminoadipic semialdehyde dehydrogenase |
| Q5JTC6 | AMER1 | APC membrane recruitment protein 1 |
| Q8IY63 | AMOTL1 | Angiomotin-like protein 1 |
| Q9H1A4 | ANAPC1 | Anaphase-promoting complex subunit 1 |
| Q9UM13 | ANAPC10 | Anaphase-promoting complex subunit 10 |
| O75179 | ANKRD17 | Ankyrin repeat domain-containing protein 17 |
| O43747 | AP1G1 | AP-1 complex subunit gamma-1 |
| Q9BXS5 | AP1M1 | AP-1 complex subunit mu-1 |
| P61966 | AP1S1 | AP-1 complex subunit sigma-1A |
| P56377 | AP1S2 | AP-1 complex subunit sigma-2 |
| O00189 | AP4M1 | AP-4 complex subunit mu-1 |
| P02649 | APOE | Apolipoprotein E |
| Q8N6T3 | ARFGAP1 | ADP-ribosylation factor GTPase-activating protein 1 |
| Q9NP61 | ARFGAP3 | ADP-ribosylation factor GTPase-activating protein 3 |
| P05089 | ARG1 | Arginase-1 |
| Q8N264 | ARHGAP24 | Rho GTPase-activating protein 24 |
| Q6ZSZ5 | ARHGEF18 | Rho guanine nucleotide exchange factor 18 |
| P25705 | ATP5F1A | ATP synthase subunit alpha, mitochondrial |
| P06576 | ATP5F1B | ATP synthase subunit beta, mitochondrial |
| P36542 | ATP5F1C | ATP synthase subunit gamma, mitochondrial |
| P56134 | ATP5MF | ATP synthase subunit f, mitochondrial |
| O75964 | ATP5MG | ATP synthase subunit g, mitochondrial |
| P24539 | ATP5PB | ATP synthase F(0) complex subunit B1, mitochondrial |
| O75947 | ATP5PD | ATP synthase subunit d, mitochondrial |
| P18859 | ATP5PF | ATP synthase-coupling factor 6, mitochondrial |
| P48047 | ATP5PO | ATP synthase subunit O, mitochondrial |
| O14965 | AURKA | Aurora kinase A |
| Q96GD4 | AURKB | Aurora kinase B |
| Q9UQB9 | AURKC | Aurora kinase C |
| P30530 | AXL | Tyrosine-protein kinase receptor UFO |
| Q99728 | BARD1 | BRCA1-associated RING domain protein 1 |
| P50895 | BCAM | Basal cell adhesion molecule |
| P56945 | BCAR1 | Breast cancer anti-estrogen resistance protein 1 |
| Q9NYF8 | BCLAF1 | Bcl-2-associated transcription factor 1 |
| P11274 | BCR | Breakpoint cluster region protein |
| P54132 | BLM | Bloom syndrome protein |
| P38398 | BRCA1 | Breast cancer type 1 susceptibility protein |
| P51587 | BRCA2 | Breast cancer type 2 susceptibility protein |
| P35613 | BSG | Basigin |
| Q07021 | C1QBP | Complement component 1 Q subcomponent-binding protein, mitochondrial |
| Q9BXJ1 | C1QTNF1 | Complement C1q tumor necrosis factor-related protein 1 |
| P27824 | CANX | Calnexin |
| P16152 | CBR1 | Carbonyl reductase [NADPH] 1 |
| O75828 | CBR3 | Carbonyl reductase [NADPH] 3 |
| O00626 | CCL22 | C-C motif chemokine 22 |
| O00626 | CCL22 | C-C motif chemokine 22 |
| P55773 | CCL23 | C-C motif chemokine 23 |
| O15444 | CCL25 | C-C motif chemokine 25 |
| P13501 | CCL5 | C-C motif chemokine 5 |
| P20248 | CCNA2 | Cyclin-A2 |
| P14635 | CCNB1 | G2/mitotic-specific cyclin-B1 |
| P51946 | CCNH | Cyclin-H |
| O60583 | CCNT2 | Cyclin-T2 |
| Q8ND76 | CCNY | Cyclin-Y |
| P41597 | CCR2 | C-C chemokine receptor type 2 |
| P51679 | CCR4 | C-C chemokine receptor type 4 |
| P51681 | CCR5 | C-C chemokine receptor type 5 |
| P32248 | CCR7 | C-C chemokine receptor type 7 |
| P32248 | CCR7 | C-C chemokine receptor type 7 |
| P78371 | CCT2 | T-complex protein 1 subunit beta |
| P08571 | CD14 | Monocyte differentiation antigen CD14 |
| Q86VB7 | CD163 | Scavenger receptor cysteine-rich type 1 protein M130 |
| P21757 | CD204/MSR1 | Macrophage scavenger receptor |
| P20138 | CD33 | Myeloid cell surface antigen CD33 |
| P20138 | CD33 | Myeloid cell surface antigen CD33 |
| P25942 | CD40 | Tumor necrosis factor receptor superfamily member 5 |
| Q07108 | CD69 | Early activation antigen CD69 |
| P33681 | CD80 | T-lymphocyte activation antigen CD80 |
| P42081 | CD86 | T-lymphocyte activation antigen CD86 |
| P19022 | CDH2 | Cadherin-2 |
| P55283 | CDH4 | Cadherin-4 |
| Q6ZTQ4 | CDHR3 | Cadherin-related family member 3 |
| P06493 | CDK1 | Cyclin-dependent kinase 1 |
| P24941 | CDK2 | Cyclin-dependent kinase 2 |
| P11802 | CDK4 | Cyclin-dependent kinase 4 |
| Q00534 | CDK6 | Cyclin-dependent kinase 6 |
| P38936 | CDKN1A | Cyclin-dependent kinase inhibitor 1 |
| P42771 | CDKN2A | Cyclin-dependent kinase inhibitor 2A |
| Q9H211 | CDT1 | DNA replication factor Cdt1 |
| P17676 | CEBPB | CCAAT/enhancer-binding protein beta |
| P49715 | CEPBA | CCAAT/enhancer-binding protein alpha |
| Q13111 | CHAF1A | Chromatin assembly factor 1 subunit A |
| Q7Z7A1 | CNTRL | Centriolin |
| P39060 | COL18A1 | Collagen alpha-1(XVIII) chain |
| P0245 | COL1A1 | Collagen alpha-1(I) chain |
| P08123 | COL1A2 | Collagen alpha-2(I) chain |
| Q8NFW1 | COL22A1 | Collagen alpha-1(XXII) chain |
| P08572 | COL4A2 | Collagen alpha-2(IV) chain |
| Q01955 | COL4A3 | Collagen alpha-3(IV) chain |
| P29400 | COL4A5 | Collagen alpha-5(IV) chain |
| Q14031 | COL4A6 | Collagen alpha-6(IV) chain |
| P20908 | COL5A1 | Collagen alpha-1(V) chain |
| P12109 | COL6A1 | Collagen alpha-1(VI) chain |
| Q02388 | COL7A1 | Collagen alpha-1(VII) chain |
| P53618 | COPB1 | Coatomer subunit beta |
| P35606 | COPB2 | Coatomer subunit beta' |
| O14579 | COPE | Coatomer subunit epsilon |
| Q9Y678 | COPG1 | Coatomer subunit gamma-1 |
| Q9Y6N1 | COX11 | Cytochrome c oxidase assembly protein COX11, mitochondrial |
| Q7KZN9 | COX15 | Cytochrome c oxidase assembly protein COX15 |
| P09669 | COX6C | Cytochrome c oxidase subunit 6C |
| Q14194 | CRMP1 | Dihydropyrimidinase-related protein 1 |
| P07333 | CSF1R | Macrophage colony-stimulating factor 1 receptor |
| P41240 | CSK | Tyrosine-protein kinase CSK |
| P35222 | CTNNB1 | Catenin beta-1 |
| P07858 | CTSB | Cathepsin B |
| P07339 | CTSD | Cathepsin D |
| P07711 | CTSL | Cathepsin L1 |
| Q13618 | CUL3 | Cullin-3 |
| P09341 | CXCL1 | Growth-regulated alpha protein |
| P48061 | CXCL12/SDF1 | Stromal cell-derived factor 1 |
| P42830 | CXCL5 | C-X-C motif chemokine 5 |
| P80162 | CXCL6 | C-X-C motif chemokine 6 |
| Q16832 | DDR2 | Discoidin domain-containing receptor 2 |
| Q9BUN8 | DERL1 | Derlin-1 |
| Q7L2E3 | DHX30 | ATP-dependent RNA helicase DHX30 |
| O75165 | DNAJC13 | DnaJ homolog subfamily C member 13 |
| Q05193 | DNM1 | Dynamin-1 |
| Q9UQ16 | DNM3 | Dynamin-3 |
| Q14185 | DOCK1 | Dedicator of cytokinesis protein 1 |
| Q16555 | DPYSL2 | Dihydropyrimidinase-related protein 2 |
| P15924 | DSP | Desmoplakin |
| Q96AV8 | E2F7 | Transcription factor E2F7 |
| P30084 | ECHS1 | Enoyl-CoA hydratase, mitochondrial |
| P42126 | ECI1 | Enoyl-CoA delta isomerase 1, mitochondrial |
| P00533 | EGFR | Epidermal growth factor receptor |
| P47813 | EIF1AX | Eukaryotic translation initiation factor 1A, X-chromosomal |
| P49770 | EIF2B2 | Translation initiation factor eIF-2B subunit beta |
| P41214 | EIF2D | Eukaryotic translation initiation factor 2D |
| Q92556 | ELMO1 | Engulfment and cell motility protein 1 |
| Q96JJ3 | ELMO2 | Engulfment and cell motility protein 2 |
| P17813 | ENG | Endoglin |
| Q9NZ08 | ERAP1 | Endoplasmic reticulum aminopeptidase 1 |
| Q6P179 | ERAP2 | Endoplasmic reticulum aminopeptidase 2 |
| P07992 | ERCC1 | DNA excision repair protein ERCC-1 |
| P19447 | ERCC3 | General transcription and DNA repair factor IIH helicase subunit XPB |
| P38117 | ETFB | Electron transfer flavoprotein subunit beta |
| Q96A65 | EXOC4 | Exocyst complex component 4 |
| Q9NQT4 | EXOSC5 | Exosome complex component RRP46 |
| Q15910 | EZH2 | Histone-lysine N-methyltransferase EZH2 |
| P35556 | FBN2 | Fibrillin-2 |
| P05230 | FGF1 | Fibroblast growth factor 1 |
| P17948 | FLT1/VEGFR1 | Vascular endothelial growth factor receptor 1 |
| P02751 | FN1 | Fibronectin |
| P04066 | FUCA1 | Tissue alpha-L-fucosidase |
| P06241 | FYN | Tyrosine-protein kinase Fyn |
| Q8TAE8 | GADD45GIP1 | Growth arrest and DNA damage-inducible proteins-interacting protein 1 |
| P04406 | GAPDH | Glyceraldehyde-3-phosphate dehydrogenase |
| P32455 | GBP1 | Guanylate-binding protein 1 |
| P14136 | GFAP | Glial fibrillary acidic protein |
| Q96RP9 | GFM1 | Elongation factor G, mitochondrial |
| Q969S9 | GFM2 | Ribosome-releasing factor 2, mitochondrial |
| Q9NZM5 | GLTSCR2 | NOP53 Ribosome Biogenesis Factor |
| P49841 | GSK3B | Glycogen synthase kinase-3 beta |
| Q972Q3 | GSTK1 | Glutathione S-transferase kappa 1 |
| P78417 | GSTO1 | Glutathione S-transferase omega-1 |
| P29083 | GTF2E1 | General transcription factor IIE subunit 1 |
| P35269 | GTF2F1 | General transcription factor IIF subunit 1 |
| P32780 | GTF2H1 | General transcription factor IIH subunit 1 |
| Q13888 | GTF2H2 | General transcription factor IIH subunit 2 |
| Q13889 | GTF2H3 | General transcription factor IIH subunit 3 |
| Q6EKJ0 | GTF2IRD2B | General transcription factor II-I repeat domain-containing protein 2B |
| Q8WUA4 | GTF3C2 | General transcription factor 3C polypeptide 2 |
| Q9Y5Q9 | GTF3C3 | General transcription factor 3C polypeptide 3 |
| Q13547 | HDAC1 | Histone deacetylase 1 |
| Q92769 | HDAC2 | Histone deacetylase 2 |
| Q9H583 | HEATR1 | HEAT repeat-containing protein 1 |
| P07686 | HEXB | Beta-hexosaminidase subunit beta |
| O94992 | HEXIM1 | Protein HEXIM1 |
| P05534 | HLA-A | HLA class I histocompatibility antigen, A-24 alpha chain |
| P30505 | HLA-C | HLA class I histocompatibility antigen, Cw-8 alpha chain |
| P30510 | HLA-C | HLA class I histocompatibility antigen, Cw-14 alpha chain |
| P06340 | HLA-DOA | HLA class II histocompatibility antigen, DO alpha chain |
| P26583 | HMGB2 | High mobility group protein B2 |
| Q99714 | HSD17B10 | 3-hydroxyacyl-CoA dehydrogenase type-2 |
| O43464 | HTRA2 | Serine protease HTRA2, mitochondrial |
| P09914 | IFIT1 | Interferon-induced protein with tetratricopeptide repeats 1 |
| P01579 | IFNG | Interferon gamma (IFN-γ) |
| Q16270 | IGFBP7 | Insulin-like growth factor-binding protein 7 |
| P01584 | IL-1β | Interleukin-1 beta |
| P29459 | IL-12 | Interleukin-12 subunit alpha |
| P05231 | IL-6 | Interleukin-6 |
| P01589 | IL2RA | Interleukin-2 receptor subunit alpha |
| Q12906 | ILF3 | Interleukin enhancer-binding factor 3 |
| Q13418 | ILK | Integrin-linked protein kinase |
| Q9ULG1 | INO80 | Chromatin-remodeling ATPase |
| P05161 | ISG15 | Ubiquitin-like protein ISG15 |
| Q96AZ6 | ISG20 | Interferon-stimulated gene 20 kDa protein |
| P26006 | ITGA3 | Integrin alpha-3 |
| P23229 | ITGA6 | Integrin alpha-6 |
| P06756 | ITGAV | Integrin alpha-V |
| P05556 | ITGB1 | Integrin beta-1 |
| P16144 | ITGB4 | Integrin beta-4 |
| P18084 | ITGB5 | Integrin beta-5 |
| P14923 | JUP | Junction plakoglobin |
| P35968 | KDR/VGFR2 | Vascular endothelial growth factor receptor 2 |
| O15091 | KIAA0391 | Mitochondrial ribonuclease P catalytic subunit |
| O00139 | KIF2A | Kinesin-like protein KIF2A |
| O95239 | KIF4A | Chromosome-associated kinesin KIF4A |
| Q2VIQ3 | KIF4B | Chromosome-associated kinesin KIF4B |
| Q9BW19 | KIFC1 | Kinesin-like protein KIF1C |
| O95198 | KLHL2 | Kelch-like protein 2 |
| Q9P2J3 | KLHL9 | Kelch-like protein 9 |
| P50748 | KNTC1 | Kinetochore-associated protein 1 |
| P32004 | L1CAM | Neural cell adhesion molecule L1 |
| P11047 | LAMC1 | Laminin subunit gamma-1 |
| Q6IAA8 | LAMTOR1 | Regulator complex protein LAMTOR1 |
| P13796 | LCP1 | Plastin-2 |
| P01130 | LDLR | Low-density lipoprotein receptor |
| Q8IVL6 | LEPREL2/P3H3 | Prolyl 3-Hydroxylase 3 |
| P49916 | LIG3 | DNA ligase 3 |
| Q9UIQ6 | LNPEP | Leucyl-cystinyl aminopeptidase |
| P28300 | LOX | Protein-lysine 6-oxidase |
| Q07954 | LRP1 | LDL Receptor Related Protein 1/Type V TGF-Beta receptor |
| Q9BRK4 | LZTS2 | Leucine zipper putative tumor suppressor 2 |
| Q02750 | MAP2K1 | Dual specificity mitogen-activated protein kinase kinase 1 |
| P36507 | MAP2K2 | Dual specificity mitogen-activated protein kinase kinase 2 |
| Q16539 | MAPK14 | Mitogen-activated protein kinase 14 |
| Q7Z434 | MAVS | Mitochondrial antiviral-signaling protein |
| P43121 | MCAM | Cell surface glycoprotein MUC18 |
| Q7L590 | MCM10 | Protein MCM10 homolog |
| P49736 | MCM2 | DNA replication licensing factor MCM2 |
| P25205 | MCM3 | DNA replication licensing factor MCM3 |
| P33991 | MCM4 | DNA replication licensing factor MCM4 |
| P33992 | MCM5 | DNA replication licensing factor MCM5 |
| Q14566 | MCM6 | DNA replication licensing factor MCM6 |
| P33993 | MCM7 | DNA replication licensing factor MCM7 |
| Q9Y316 | MEMO1 | Protein MEMO1 |
| O00255 | MEN1 | Menin |
| Q12866 | MERTK | Tyrosine-protein kinase Mer |
| Q12866 | MERTK | Tyrosine-protein kinase Mer |
| P08581 | MET | Hepatocyte growth factor receptor |
| P46013 | MKI67 | Proliferation marker protein Ki-67 |
| P03886 | MT-ND1 | NADH-ubiquinone oxidoreductase chain 1 |
| P03915 | MT-ND5 | NADH-ubiquinone oxidoreductase chain 5 |
| Q9NZJ7 | MTCH1 | Mitochondrial carrier homolog 1 |
| P42345 | MTOR | Serine/threonine-protein kinase mTOR |
| P20591 | MX1 | Interferon-induced GTP-binding protein Mx1 |
| P20592 | MX2 | Interferon-induced GTP-binding protein Mx2 |
| P10242 | MYB | Transcriptional activator Myb |
| Q99836 | MYD88 | Myeloid differentiation primary response protein MyD88 |
| P54920 | NAPA | Alpha-soluble NSF attachment protein |
| Q9HC98 | NEK6 | Serine/threonine-protein kinase Nek6 |
| Q99519 | NEU1 | Sialidase-1 |
| O00712 | NFIB | Nuclear factor 1 B-type |
| P19838 | NFKB1 | Nuclear factor NF-kappa-B |
| O00221 | NFKBIE | NF-kappa-B inhibitor epsilon |
| P01138 | NGF | Beta-nerve growth factor |
| Q8N5F7 | NKAP | NF-kappa-B-activating protein |
| Q9NPP4 | NLRC4 | NLR family CARD domain-containing protein 4 |
| Q9Y239 | NOD1 | Nucleotide-binding oligomerization domain-containing protein 1 |
| Q9HC29 | NOD2 | Nucleotide-binding oligomerization domain-containing protein 2 |
| Q9H8H0 | NOL11 | Nucleolar protein 11 |
| Q14978 | NOLC1 | Nucleolar and coiled-body phosphoprotein 1 |
| P78316 | NOP14 | Nucleolar protein 14 |
| P55786 | NPEPPS | Puromycin-sensitive aminopeptidase |
| P01111 | NRAS | GTPase NRas |
| O95478 | NSA2 | Ribosome biogenesis protein NSA2 homolog |
| Q96PB7 | OLFM3 | Noelin-3 |
| Q9UBD5 | ORC3 | Origin recognition complex subunit 3 |
| O43929 | ORC4 | Origin recognition complex subunit 4 |
| Q9H244 | P2RY12 | P2Y purinoceptor 12 |
| Q32P28 | P3H1 | Prolyl 3-hydroxylase 1 |
| Q8IVL6 | P3H3 | Prolyl 3-hydroxylase 3 |
| O15460 | P4HA2 | Prolyl 4-hydroxylase subunit alpha-2 |
| Q13153 | PAK1 | Serine/threonine-protein kinase PAK 1 |
| Q8WX93 | PALLD | Palladin |
| P09619 | PDGFRB | Platelet-derived growth factor receptor beta |
| Q15118 | PDK1 | [Pyruvate dehydrogenase (acetyl-transferring)] kinase isozyme 1, mitochondrial |
| P30086 | PEBP1 | Phosphatidylethanolamine-binding protein 1 |
| P35232 | PHB | Prohibitin |
| P42338 | PIK3CB | Phosphatidylinositol 4,5-bisphosphate 3-kinase catalytic subunit beta isoform |
| P53350 | PLK1 | Serine/threonine-protein kinase PLK1 |
| P29590 | PML | Protein PML |
| Q8TCS8 | PNPT1 | Polyribonucleotide nucleotidyltransferase 1, mitochondrial |
| P49005 | POLD2 | DNA polymerase delta subunit 2 |
| Q07864 | POLE | DNA polymerase epsilon catalytic subunit A |
| Q9UBT6 | POLK | DNA polymerase kappa |
| P24928 | POLR2A | DNA-directed RNA polymerase II subunit RPB1 |
| P19387 | POLR2C | DNA-directed RNA polymerase II subunit RPB3 |
| P62487 | POLR2G | DNA-directed RNA polymerase II subunit RPB7 |
| P52435 | POLR2J | DNA-directed RNA polymerase II subunit RPB11-a |
| Q15173 | PPP2R5B | Serine/threonine-protein phosphatase 2A 56 kDa regulatory subunit beta isoform |
| O60828 | PQBP1 | Polyglutamine-binding protein 1 |
| P30048 | PRDX3 | Thioredoxin-dependent peroxide reductase, mitochondrial |
| Q13162 | PRDX4 | Peroxiredoxin-4 |
| P30044 | PRDX5 | Peroxiredoxin-5 |
| P30041 | PRDX6 | Peroxiredoxin-6 |
| P49642 | PRIM1 | DNA primase small subunit |
| P07225 | PROS1 | Vitamin K-dependent protein S |
| P40306 | PSMB10 | Proteasome subunit beta type-10 |
| P28074 | PSMB5 | Proteasome subunit beta type-5 |
| P28072 | PSMB6 | Proteasome subunit beta type-6 |
| Q99436 | PSMB7 | Proteasome subunit beta type-7 |
| P28062 | PSMB8 | Proteasome subunit beta type-8 |
| P28065 | PSMB9 | Proteasome subunit beta type-9 |
| Q06323 | PSME1 | Proteasome activator complex subunit 1 |
| Q9UL46 | PSME2 | Proteasome activator complex subunit 2 |
| P61289 | PSME3 | Proteasome activator complex subunit 3 |
| Q13308 | PTK7 | Inactive tyrosine-protein kinase 7 |
| Q06124 | PTPN11 | Tyrosine-protein phosphatase non-receptor type 11 |
| P08575 | PTPRC | Receptor-type tyrosine-protein phosphatase C |
| P23470 | PTPRG | Receptor-type tyrosine-protein phosphatase gamma |
| Q92626 | PXDN | Peroxidasin |
| P51153 | RAB13 | Ras-related protein Rab-13 |
| P20338 | RAB4A | Ras-related protein Rab-4A |
| P20339 | RAB5A | Ras-related protein Rab-5A |
| Q9H0H5 | RACGAP1 | Rac GTPase-activating protein 1 |
| Q06609 | RAD51 | DNA repair protein RAD51 homolog 1 |
| P10114 | RAP2A | Ras-related protein Rap-2a |
| Q9Y3L5 | RAP2C | Ras-related protein Rap-2c |
| Q13905 | RAPGEF1 | Rap guanine nucleotide exchange factor 1 |
| Q04864 | REL | Proto-oncogene c-Rel |
| Q04206 | RELA | Transcription factor p65 |
| P35251 | RFC1 | Replication factor C subunit 1 |
| P40938 | RFC3 | Replication factor C subunit 3 |
| P40937 | RFC5 | Replication factor C subunit 5 |
| P61586 | RHOA | Transforming protein RhoA |
| Q63HN8 | RNF213 | E3 ubiquitin-protein ligase RNF213 |
| Q13464 | ROCK1 | Rho-associated protein kinase 1 |
| O75582 | RPS6KA5 | Ribosomal protein S6 kinase alpha-5 |
| Q8N122 | RPTOR | Regulatory-associated protein of mTOR |
| P10301 | RRAS | Ras-related protein R-Ras |
| Q15050 | RRS1 | Ribosome biogenesis regulatory protein homolog |
| Q9UHA3 | RSL24D1 | Probable ribosome biogenesis protein RLP24 |
| Q9NSC2 | SALL1 | Sal-like protein 1 |
| Q9Y3Z3 | SAMHD1 | Deoxynucleoside triphosphate triphosphohydrolase |
| P31040 | SDHA | Succinate dehydrogenase [ubiquinone] flavoprotein subunit, mitochondrial |
| P21912 | SDHB | Succinate dehydrogenase [ubiquinone] iron-sulfur subunit, mitochondrial |
| P05121 | SERPINE1 | Plasminogen activator inhibitor 1 |
| Q9Y371 | SH3GLB1 | Endophilin-B1 |
| P29353 | SHC1 | SHC-transforming protein 1 |
| Q13309 | SKP2 | S-phase kinase-associated protein 2 |
| Q15796 | SMAD2 | Mothers against decapentaplegic homolog 2 |
| P84022 | SMAD3 | Mothers against decapentaplegic homolog 3 |
| Q13485 | SMAD4 | Mothers against decapentaplegic homolog 4 |
| Q8TAD8 | SNIP1 | Smad nuclear-interacting protein 1 |
| Q13573 | SNW1 | SNW domain-containing protein 1 |
| P04179 | SOD2 | Superoxide dismutase [Mn], mitochondrial |
| P23497 | SP100 | Nuclear autoantigen Sp-100 |
| P09486 | SPARC | Secreted Protein Acidic And Cysteine Rich |
| P12931 | SRC | Proto-oncogene tyrosine-protein kinase Src |
| P42224 | STAT1 | Signal transducer and activator of transcription 1 |
| P52630 | STAT2 | Signal transducer and activator of transcription 2 |
| P40763 | STAT3 | Signal transducer and activator of transcription 3 |
| P42226 | STAT6 | Signal transducer and activator of transcription 6 |
| Q9Y6E0 | STK24 | Serine/threonine-protein kinase 24 |
| Q9P289 | STK26/MST4 | Serine/threonine-protein kinase 26 |
| Q86Y82 | STX12 | Syntaxin-12 |
| Q15750 | TAB1 | TGF-beta-activated kinase 1 and MAP3K7-binding protein 1 |
| Q9BW92 | TARS2 | Threonine--tRNA ligase, mitochondrial |
| P17987 | TCP1 | T-complex protein 1 subunit alpha |
| Q00059 | TFAM | Transcription factor A, mitochondrial |
| Q9H5Q4 | TFB2M | Dimethyladenosine transferase 2, mitochondrial |
| P02786 | TFRC | Transferrin receptor protein 1 |
| O43294 | TGFB1I1 | Transforming growth factor beta-1-induced transcript 1 protein |
| Q15582 | TGFBI | Transforming growth factor-beta-induced protein ig-h3 |
| P37173 | TGFBR2 | TGF-beta receptor type-2 |
| P07996 | THBS1 | Thrombospondin-1 |
| Q9Y2W1 | THRAP3 | Thyroid hormone receptor-associated protein 3 |
| P62072 | TIMM10 | Mitochondrial import inner membrane translocase subunit Tim10 |
| Q86UE8 | TLK2 | Serine/threonine-protein kinase tousled-like 2 |
| Q9Y490 | TLN1 | Talin-1 |
| Q9Y4G6 | TLN2 | Talin-2 |
| P49755 | TMED10 | Transmembrane emp24 domain-containing protein 10 |
| Q9BVK6 | TMED9 | Transmembrane emp24 domain-containing protein 9 |
| P82094 | TMF1 | TATA element modulatory factor |
| P24821 | TNC | Tenascin |
| P01375 | TNF-α | Tumor necrosis factor |
| Q15388 | TOMM20 | Mitochondrial import receptor subunit TOM20 |
| Q9NS69 | TOMM22 | Mitochondrial import receptor subunit TOM22 |
| O96008 | TOMM40 | Mitochondrial import receptor subunit TOM40 |
| Q9P0U1 | TOMM7 | Mitochondrial import receptor subunit TOM7 |
| Q92547 | TOPBP1 | DNA topoisomerase 2-binding protein 1 |
| Q96S44 | TP53RK | EKC/KEOPS complex subunit TP53RK |
| Q9ULW0 | TPX2 | Targeting protein for Xklp2 |
| Q13114 | TRAF3 | TNF receptor-associated factor 3 |
| Q9NZC2 | TREM2 | Triggering receptor expressed on myeloid cells 2 |
| Q8IYM9 | TRIM22 | E3 ubiquitin-protein ligase TRIM22 |
| Q14258 | TRIM25 | E3 ubiquitin/ISG15 ligase TRIM25 |
| P14373 | TRIM27 | Zinc finger protein RFP |
| Q13049 | TRIM32 | E3 ubiquitin-protein ligase TRIM32 |
| Q9BRZ2 | TRIM56 | E3 ubiquitin-protein ligase TRIM56 |
| Q8IWR1 | TRIM59 | Tripartite motif-containing protein 59 |
| Q96Q11 | TRNT1 | CCA tRNA nucleotidyltransferase 1, mitochondrial |
| Q99816 | TSG101 | Tumor susceptibility gene 101 protein |
| Q96RT7 | TUBGCP6 | Gamma-tubulin complex component 6 |
| P49411 | TUFM | Elongation factor Tu, mitochondrial |
| P61086 | UBE2K | Ubiquitin-conjugating enzyme E2 K |
| Q9H832 | UBE2Z | Ubiquitin-conjugating enzyme E2 Z |
| Q05086 | UBE3A | Ubiquitin-protein ligase E3A |
| O94874 | UFL1 | E3 UFM1-protein ligase 1 |
| Q9BVJ6 | UTP14A | U3 small nucleolar RNA-associated protein 14 homolog A |
| Q8TED0 | UTP15 | U3 small nucleolar RNA-associated protein 15 homolog |
| Q9Y5J1 | UTP18 | U3 small nucleolar RNA-associated protein 18 homolog |
| Q9NYH9 | UTP6 | U3 small nucleolar RNA-associated protein 6 homolog |
| P19320 | VCAM1 | Vascular cell adhesion protein 1 |
| P18206 | VCL | Vinculin |
| P04004 | VTN | Vitronectin |
| Q9GZL7 | WDR12 | Ribosome biogenesis protein WDR12 |
| P0C1S8 | WEE2 | Wee1-like protein kinase 2 |
| Q9H0M0 | WWP1 | NEDD4-like E3 ubiquitin-protein ligase |
| Q9Y2Z4 | YARS | Tyrosine--tRNA ligase, mitochondrial |
| O15498 | YKT6 | Synaptobrevin homolog YKT6 |
| O95405 | ZFYVE9 | Zinc finger FYVE domain-containing protein 9 |
| Q9H900 | ZWILCH | Protein zwilch homolog |
